# Shared experimental materials inflate and reorder cross-participant decoding benchmarks

**DOI:** 10.64898/2026.10.07.757228

**Authors:** Han Cui, Yuge Zhu, Shaojia Huang, Tao Xu

**Affiliations:** Department of Biomedical Engineering, College of Engineering, Shantou University, Shantou, China; School of Intelligent Manufacturing and Aeronautics, Zhuhai College of Science and Technology, Zhuhai, China

## Abstract

Decoding mental states from brain activity underpins brain–computer interfaces and cognitive neuroscience. Decoders are judged by how well they generalize to people left out of training, taken as evidence that they capture what transfers between brains. However, most benchmarks present the same experimental materials to every participant, so held-out participants are tested on materials the model has already seen. Here we show, across 15 EEG, fMRI and eye-tracking datasets and 318 matched comparisons, that shared materials inflate cross-participant accuracy by up to 46 percentage points. They also reorder methods, reversing the advantage of foundation models over handcrafted features on the largest emotion EEG benchmark. The inflation tracks cross-participant item identifiability and, in sealed prospective tests, vanished where items could not be identified. Adding training items improved new-material accuracy more reliably than adding participants. These findings call for reporting new-material accuracy alongside conventional scores and for measuring item identifiability first.

---

Translating neural activity into behavioural or cognitive states is central to understanding brain function and to neurotechnology. Implanted decoders now restore speech to people with paralysis^1,2^, and non-invasive recordings allow perceived speech to be decoded across large cohorts^3^. Yet differences between individuals and changes in recording conditions over time still limit how well models perform outside the laboratory^4,5^. Judging progress, and moving decoders towards clinical use, therefore depends on knowing whether a model can apply what it has learned under new conditions.

In recent years, many methods have been developed to predict mental states from neural signals. They range from handcrafted spectral features^6^ and compact convolutional networks such as EEGNet^7^ to self-supervised foundation models, including LaBraM, CBraMod, BIOT and EEGPT^8–11^. On a downstream task, these models adapt their representations to labelled training data through linear readouts or fine-tuning^12^, and their generalization is usually measured by predicting labels for held-out participants^8,9^. The question extends beyond EEG. Decoders of perceived speech are trained on magnetoencephalography and EEG pooled across many people^3^, and methods that align functional magnetic resonance imaging (fMRI) or electrocorticography activity between individuals usually require that all of them experienced the same materials^5,13^. Models of eye movements, behaviour and human annotation are built in the same way, from many people responding to one set of materials^14^, and their generalization is likewise judged on people left out of training. A benchmark score, however, reflects more than the model. It also depends on what the held-out recordings share with the training pool: the participants who produced them and the experimental materials that evoked them.

Testing on participants withheld from training has become the standard benchmark for generalization to new individuals^8,9^. Because most datasets present the same experimental materials to every participant, though, a held-out participant is evaluated on materials that the model has already encountered through other people. Psychology has long treated this as the fixed-effect fallacy: findings established on one set of materials need not hold for new ones^15–17^. Visual neuroscience likewise asked decoders to classify images never shown during training^18^, and related forms of data leakage inflate predictive performance in neuroimaging and genomics^19–21^. Earlier EEG studies noted that reusing the same materials raises classification scores^22–24^. What remains unknown is whether shared materials add a uniform offset or change which method appears better, and what determines the size of the effect.

Here we pair each conventional evaluation of neural decoders with one that withholds the test materials as well as the test participants (Fig. 1a). The analysis moves from the size of the resulting bias to its consequences, its mechanism, a prospective test and guidance for practice (Fig. 1b). We apply it to 15 datasets spanning EEG, fMRI and eye tracking, with 318 matched comparisons (Fig. 1c). Material reuse inflated cross-participant accuracy by up to 46 percentage points and changed model comparisons; on the largest emotion EEG benchmark, withholding materials reversed the advantage of EEG foundation models over handcrafted features. Architecture-matched controls and cross-participant item decoding show where the inflation comes from. Shared materials reward recognition of item-specific responses evoked by the materials, and the inflation tracks how well items can be identified across participants. Measured before any decoder was trained, this identifiability correctly predicted the absence of bias in sealed prospective tests. Finally, adding training items improved generalization more reliably than adding participants, which gives concrete guidance for benchmark design and data collection.

**Fig. 1.**
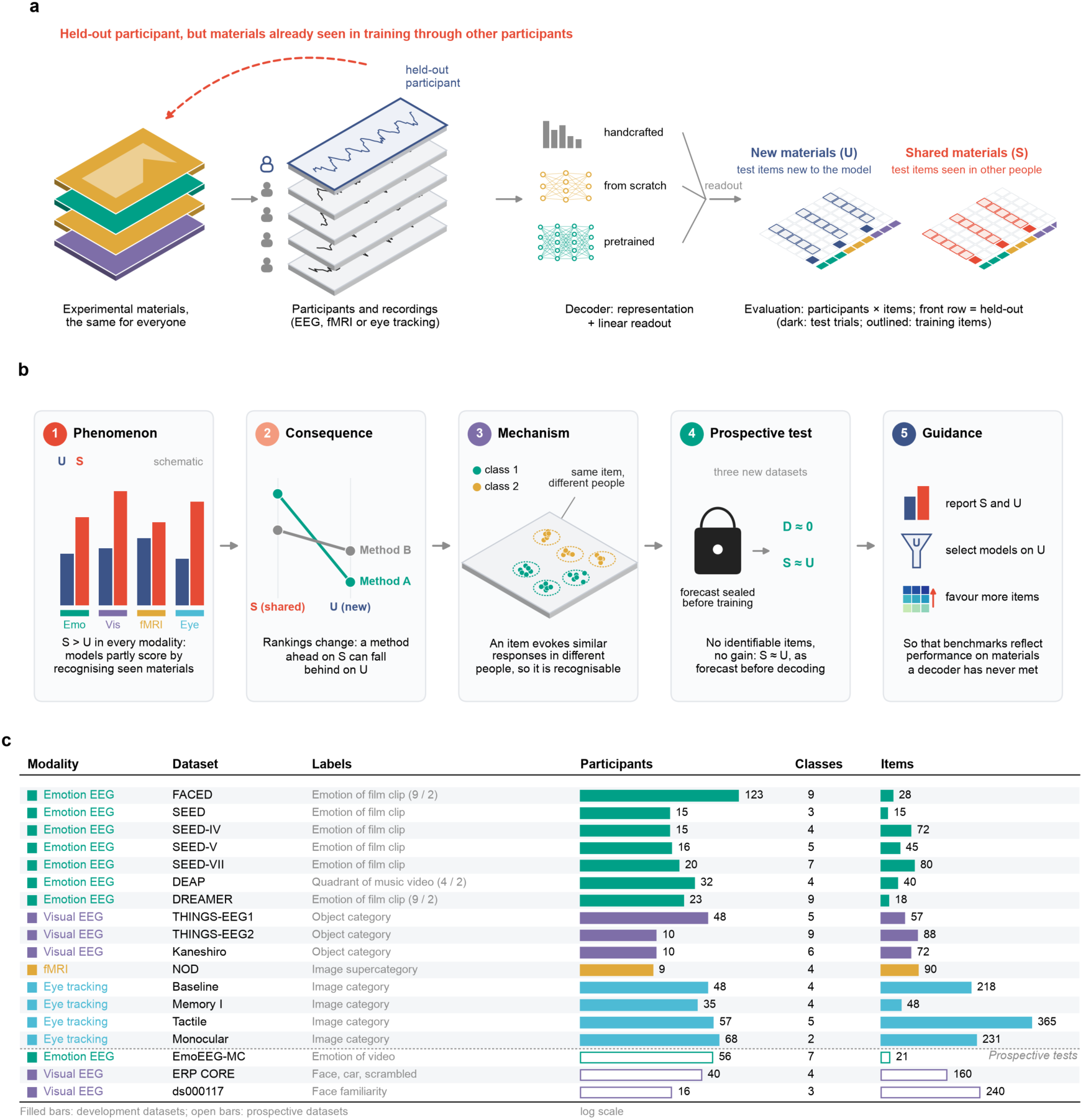
Shared experimental materials let a cross-participant test reward recognition of the materials. a, In a standard cross-participant test, the held-out participant is new to the decoder, but the materials are not, because other training participants experienced them. The paired framework scores the same test trials after training without the test materials (new-material evaluation, U) or with them (shared-material evaluation, S), with the same training participants and matched numbers of items per class and repetitions. Their difference, S − U, is the accuracy gained from having seen the test materials in other people. b, The argument of the study, from the phenomenon to its consequences, its mechanism, a prospective test and guidance for practice. D, item identifiability (Fig. 4a). c, Data sources by recording modality, with the label of each trial and the numbers of participants, classes and items. Numbers in parentheses give the classes of two tasks derived from one dataset. The four eye-tracking studies come from one database and count as one of the 15 datasets. Filled bars mark the datasets used to develop the account, and open bars the three datasets reserved for the prospective tests.

## Results

### A paired framework separates transfer to new people from transfer to new materials

Cross-participant decoding studies usually hold out participants but show every participant the same experimental materials. As a result, the individual clips, images or scenes in the test set, which we call items, have also been seen during training, through other participants. We call this shared-material evaluation (S). To isolate what such a test rewards, we paired it with new-material evaluation (U), in which neither the test participants nor the test materials appear in training (Fig. 1a). Both evaluations use the same test trials, the same training participants and matched numbers of items and repetitions per class. Their difference, the shared-material gain S − U, measures the accuracy a decoder gains from having seen the test materials in other people.

We applied this design to every data source that met three conditions: all participants experienced the same materials, class labels were fixed by the materials, and each class contained several items. The collection spans ten emotion EEG tasks from seven datasets, visual object EEG, fMRI and eye tracking (Fig. 1c). Participants and items are both random factors here, so we report balanced accuracy with intervals from a crossed bootstrap that resamples both (Methods).

### Shared materials inflate cross-participant accuracy across recording modalities

We first asked how much S exceeds U when nothing else differs. In emotion EEG, we observed shared-material gains, averaged over representations, from about 1 percentage point on DEAP^25^ to 21 points on the nine-class FACED task^26^. The gain was not limited to affective labels. Object categories decoded from visual EEG gained 11 to 12 points on THINGS-EEG1^27^ and the Kaneshiro dataset^28^, and image categories decoded from fMRI responses gained 6 points^29^ (Fig. 2a,b and Extended Data Figs. 1 and 2).

Eye movements behaved the same way. Classifying image categories from eye-movement patterns in four free-viewing studies^30^ gained between 4 and 46 points, depending on the study and the training condition (Fig. 2c and Extended Data Fig. 3a). We then checked whether anything other than the materials could explain the gain in emotion EEG. Clips cut from the same source film, presentation position, felt intensity, normalization and readout produced no consistent gain across tasks. On DREAMER, the gain shrank when labels came from each participant’s own ratings (Extended Data Fig. 4). A shared-material test thus measures two abilities at once, transfer to new people and recognition of materials the model has already met, in proportions that differ from source to source.

**Fig. 2.**
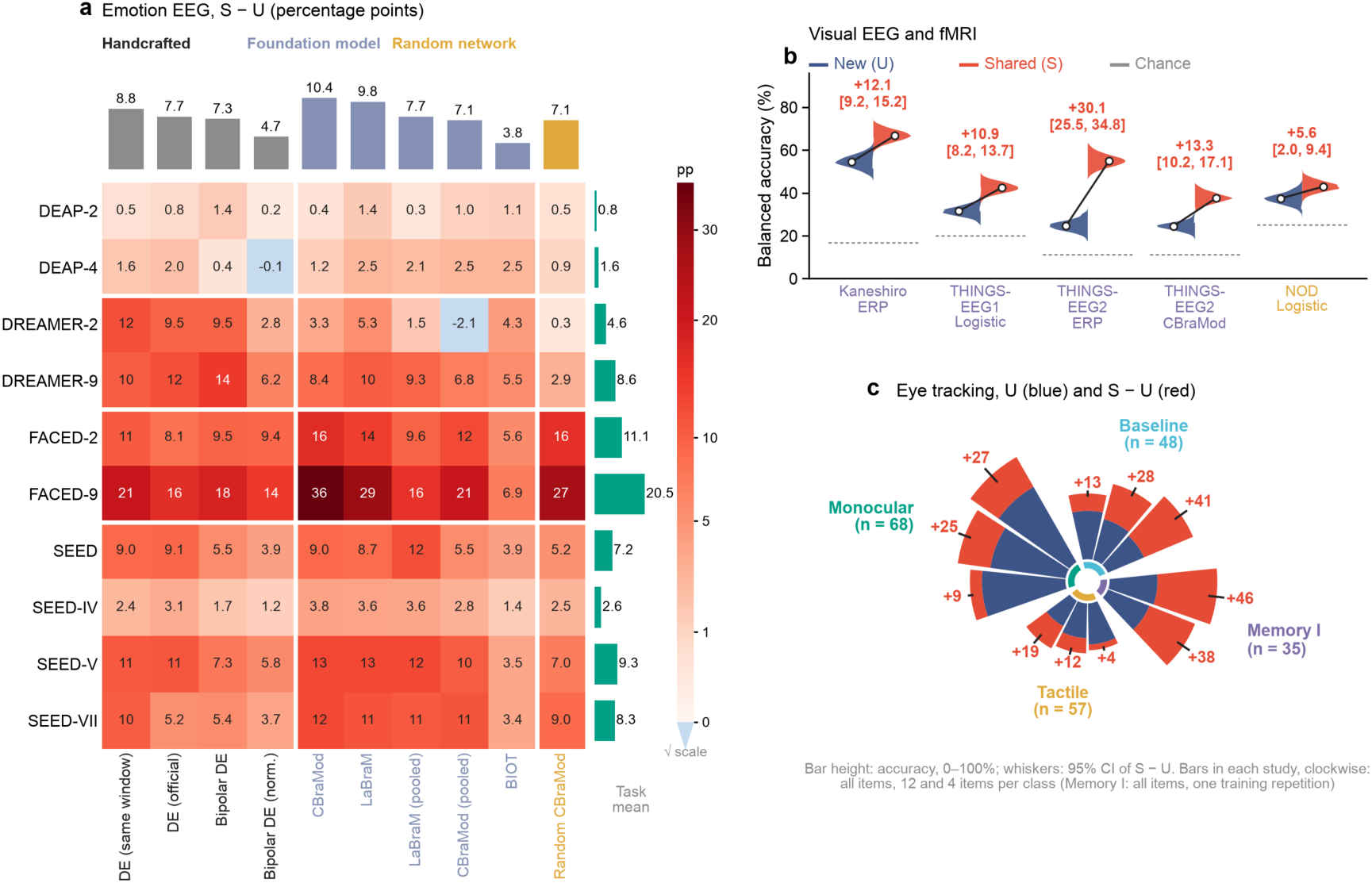
Shared materials inflate cross-participant accuracy across recording modalities. a, Shared-material gain S − U, in percentage points, for ten emotion EEG tasks (rows) and ten representations (columns); colour follows a square-root scale, and the marginal bars give the mean of each row and column. b, Balanced accuracy under S (red) and U (blue) for visual EEG and fMRI. Split violins show the bootstrap distributions, and labels give the mean gain with its 95% confidence interval. c, Accuracy under U (blue bars) and the shared-material gain (red bars) in four eye-tracking studies, after training on all items and on 12 or 4 items per class, or on all items and one repetition for Memory I. Whiskers give the 95% confidence interval of S − U. Intervals come from 2,000 crossed-bootstrap draws over participants and items; numbers of participants and items are given in Fig. 1c.

### The evaluation target changes which method appears better

A uniform offset between S and U would leave method comparisons intact. To test whether it does, we measured the advantage of one representation over another under shared materials (ΔS) and under new materials (ΔU). Their difference, the evaluation interaction I = ΔS − ΔU, is zero when the evaluation does not matter.

On FACED, every arm used two clips per class, the same training participants and the same number of windows. Under shared materials, frozen CBraMod features exceeded differential-entropy features computed on the same windows by 13 points. Under new materials, by contrast, the same comparison favoured the handcrafted features by 3 points, an interaction of 15.8 points (95% CI 12.6 to 18.9; Fig. 3a). LaBraM reversed in the same way. For both foundation models, the benchmark and the new materials gave opposite verdicts. Nor did fine-tuning remove the reversal: after linear probing and full fine-tuning, both models still led same-window differential-entropy features on FACED-9 under S and trailed them under U (Extended Data Fig. 5a and Supplementary Note 1).

This reversal was the extreme of a broader pattern. On THINGS-EEG2^31^, a baseline that reads out the event-related potential (ERP) itself outperformed CBraMod by 17 points under S but by less than 1 point under U. Similarly, the advantage of a logistic readout over a nearest-centroid readout roughly halved on THINGS-EEG1 and also shrank in fMRI, although the fMRI interaction interval included zero (Fig. 3a). Rankings moved as well. In five of the ten emotion tasks, the representation ranked first by the conventional benchmark, which trains on all items, was not the one ranked first on new materials (Fig. 3b). In short, shared-material benchmarks enlarge method differences in some sources and reorder them in others (Extended Data Fig. 6 and Supplementary Table 3), which makes them an unreliable guide to performance on new materials.

**Fig. 3.**
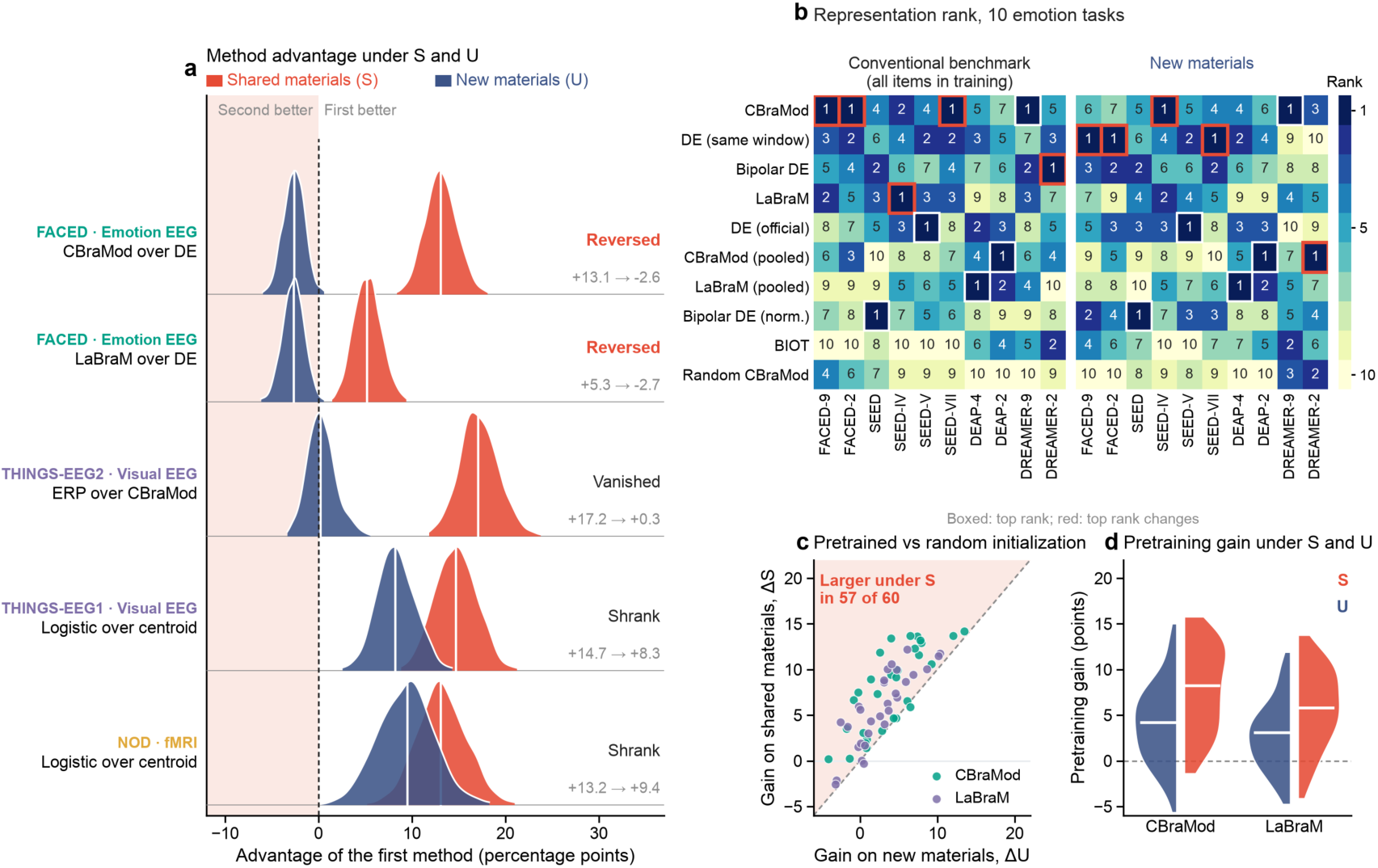
Shared-material benchmarks change which method appears better. a, Bootstrap distributions of the advantage of one method over another under S (red) and U (blue) for five comparisons; labels give the mean advantage under S and under U and whether it reversed, vanished or shrank on new materials. The FACED comparisons use the budget-matched design (n = 123 participants, 28 test clips). b, Rank of ten representations in ten emotion tasks under the conventional benchmark, which trains on all items, and under U. Boxes mark the top-ranked representation in each task and are red where it changes between the two evaluations. c, Pretraining gain, the advantage of a pretrained network over a randomly initialized network of the same architecture, under S against U for 60 comparisons, CBraMod in teal and LaBraM in purple. The dashed line marks equal gains, and the shaded region gains larger under S. d, Distributions of the same pretraining gains under S and U; white lines mark medians. Distributions in a come from 2,000 crossed-bootstrap draws.

### Architecture-matched controls separate pretraining from material recognition

Does the reversal on FACED mean that pretraining brings nothing to new materials? Comparisons with handcrafted features cannot answer this, because they change the architecture, the input and the pretraining at once. Instead, we compared each foundation model with the same architecture trained from random initialization, under an identical downstream protocol.

A randomly initialized CBraMod, read out exactly as the pretrained network, matched same-window differential-entropy features on FACED under shared materials but trailed them under new materials. The architecture alone added 6 points of shared-material gain (95% CI 2.6 to 9.6). Part of the apparent advantage of large models on standard benchmarks therefore needs no pretraining at all.

Pretraining still helped on new materials. After nested tuning, we found that CBraMod and LaBraM exceeded their randomly initialized counterparts on FACED by about 3 points each (Extended Data Fig. 5b and Supplementary Note 1). Yet the pretraining gain was larger under S than under U in 57 of the 60 comparisons of frozen pretrained and randomly initialized networks (Fig. 3c,d and Extended Data Fig. 5c). Conventional comparisons overstate what foundation models add over handcrafted features. The benefit of pretraining on new materials is real but smaller, and it becomes visible only when the architecture is held fixed.

### Cross-participant item identifiability explains the bias

A decoder can profit from shared materials only if the response to a given item is similar enough across people to be recognized in a new participant. We quantified this property as item identifiability D, the chance-corrected accuracy with which a linear probe tells apart individual items of the same class in held-out participants (Fig. 4a). Across 100 pairs of representation and task in emotion EEG, D tracked the shared-material gain closely, with a Spearman correlation of 0.87 that held within tasks (Fig. 4b and Extended Data Fig. 7; Supplementary Table 2).

If identifiability drives the bias, the representation with the higher D should receive the larger shared-material advantage. We tested this on six fixed representation pairs from FACED and THINGS-EEG2, with probes trained only on the participants and items available to the new-material arm. In all five pairs whose difference in identifiability was clearly non-zero, its sign matched the sign of the evaluation interaction (Fig. 4c). CBraMod, for example, identified FACED clips better than differential-entropy features and received the larger shared-material advantage. On THINGS-EEG2 the order flipped. There, the ERP baseline identified images better and was the representation favoured under S.

Next, we disrupted the proposed signal directly. Projecting the item-identity directions out of the frozen FACED-9 features reduced the shared-material gain of CBraMod and LaBraM by about 5 points more than removing random directions of the same rank, while new-material accuracy stayed within 1 point of its original value (Fig. 4d). The reduction was smaller on FACED-2, the valence task of FACED. On SEED-V, which has nine clips per class, the item-identity directions also carried emotion information, so the projection lowered new-material accuracy as well.

We then asked whether the identity signal came from the eyes. It was not confined to the channels nearest them. Removing the eight frontopolar, lateral frontal, temporal and mastoid channels most exposed to ocular artefacts lowered D by about one tenth, and the posterior channels alone retained about 78% of the full-montage identifiability (Fig. 4e and Extended Data Fig. 8). Measured one electrode at a time, identifiability peaked over lateral frontotemporal sites, where horizontal eye movements also project; even so, each occipital electrode retained about 60% of that peak (Extended Data Fig. 8b). Shared-material inflation, then, follows from EEG responses that are evoked by the materials, shared across people and learnable by a decoder (Extended Data Fig. 9).

### Where items cannot be identified, the bias disappears

This account makes a falsifiable prediction: where neither representation can identify individual items, S and U should agree. We tested it in three EEG datasets that had not entered any earlier analysis. For each dataset, we fixed the data definition, the probe, the forecasting rule and the decision criteria, and sealed the forecast before any category decoder was trained (Methods).

In ERP CORE^32^ and EmoEEG-MC^33^, identifiability was close to zero for both representations, so the sealed directional rule abstained. As expected, both datasets showed shared-material gains of 3.4 points or less (Fig. 5a). For the face-recognition dataset ds000117^34^, we sealed an explicit null prediction: if neither representation reached an identifiability of 0.05, the shared-material gain of each representation and their interaction would all lie within ±3 points. Identifiability stayed below 0.01 for both ERP and CBraMod features. The prediction held for all three endpoints (Fig. 5c), with an interaction of −0.7 points (90% CI −2.0 to 0.6; Extended Data Fig. 10a).

When we placed the development and prospective datasets together, they formed a single gradient (Fig. 5b). Gains stayed within a few points where identifiability was near zero and reached 13 to 37 points where it exceeded 0.3. Whether a shared-material benchmark overstates generalization can thus be judged before any decoder is compared, simply by measuring how well individual items can be identified across participants.

**Fig. 4.**
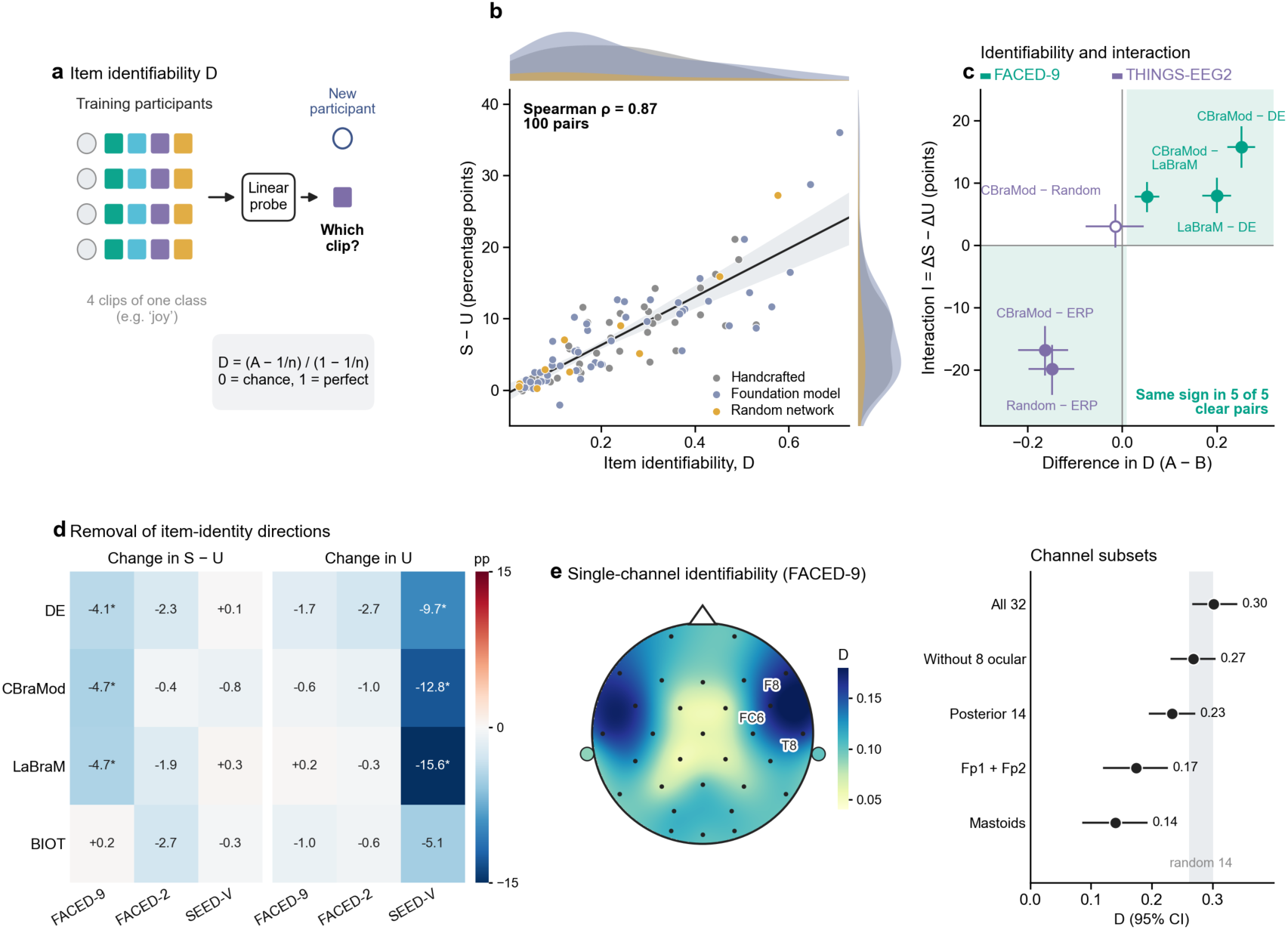
Cross-participant item identifiability explains the bias. a, Item identifiability D is the chance-corrected accuracy with which a linear probe, trained on some participants, tells apart items of the same class in held-out participants. b, D against S − U for 100 representation–task pairs in emotion EEG, coloured by representation type, with a linear fit and its 95% bootstrap band; ρ, Spearman correlation. c, Difference in D between two representations against their evaluation interaction I = ΔS − ΔU for six pairs; the open symbol marks the pair whose difference in D did not exclude zero. d, Change in S − U and in U after projecting out item-identity directions, relative to removing the same number of random directions, for four representations in three tasks; asterisks mark 95% intervals that exclude zero. e, Single-channel D for same-window DE on FACED-9, with the three highest electrodes labelled, and D of channel subsets; the grey band spans ten random 14-channel subsets. Here D is estimated with participant halves (Methods), so its values are not directly comparable with those in b. Error bars and intervals are 95% confidence intervals from 2,000 crossed-bootstrap draws.

### Consequences for data collection and model selection

If the bias depends on how well items can be recognized, design choices that make each item’s responses more reproducible should inflate S more than U. Repeated presentation is one such choice. On THINGS-EEG2, when we used all training repetitions instead of four, S rose by 28 points but U by only 3 points. The Kaneshiro dataset showed the same asymmetry on a smaller scale (Fig. 6a and Extended Data Fig. 2).

What does improve accuracy on new materials? We kept the test set fixed and doubled either the training items per class or the training participants. More items raised U more than more participants in three of the eleven sources, and more participants was clearly better in none (Fig. 6b,c and Extended Data Fig. 3b,c). The clearest case was Monocular, the eye-tracking study with the most participants, where more items added about twice as much accuracy as more participants. Items also led on FACED-9 and SEED-V. In the remaining sources, the two gains could not be told apart. For decoders meant to generalize beyond a fixed set of materials, material diversity is the safer investment.

**Fig. 5.**
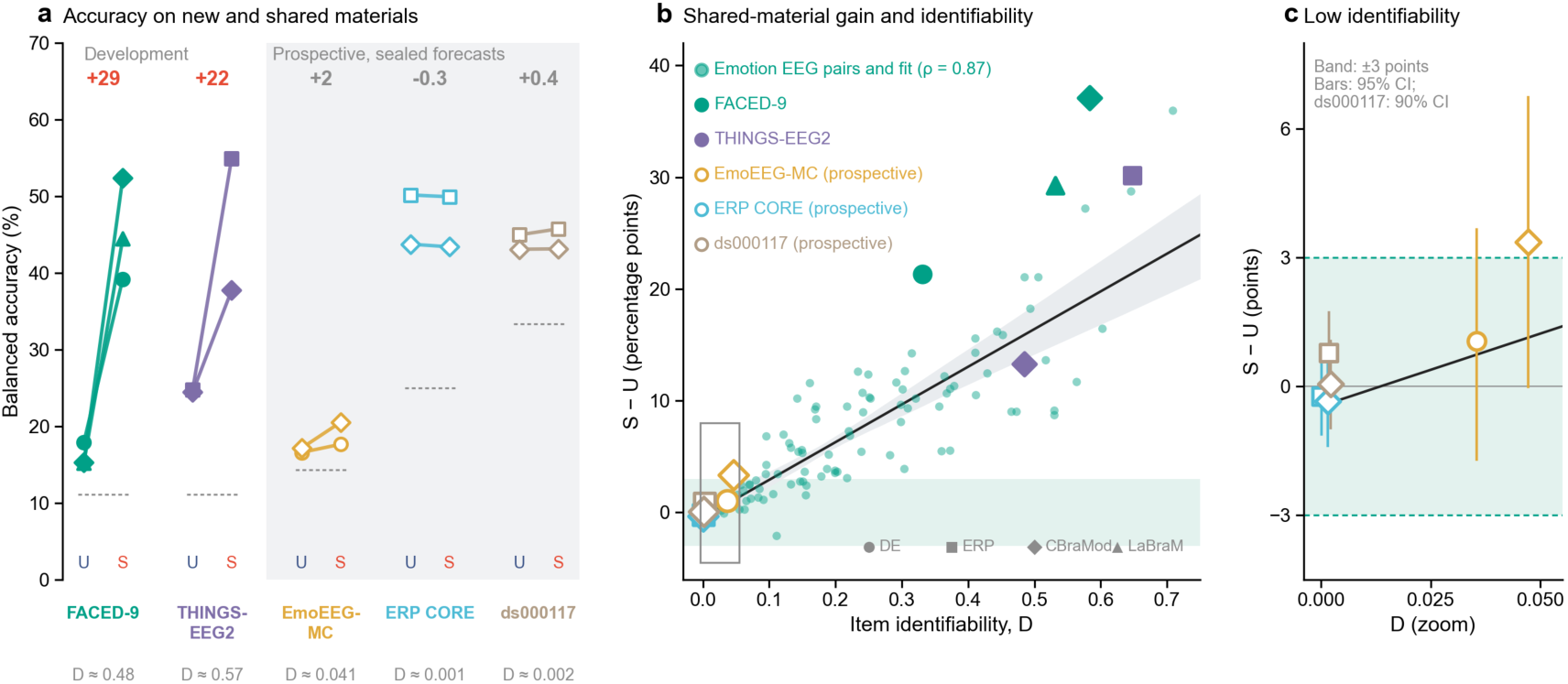
Where items cannot be identified, the bias disappears. a, Balanced accuracy under U and S for each representation in two development datasets and three prospective datasets. Numbers above each dataset give its mean shared-material gain across the representations shown, and values below its mean item identifiability D. b, Shared-material gain against D for the 100 emotion representation–task pairs (small dots) and for the datasets in a (filled symbols, development; open symbols, prospective), with a linear fit and its 95% bootstrap band; the shaded band marks ±3 points. For the datasets in a, D was estimated within the new-material training pools and FACED-9 used the budget-matched design (Methods), so these symbols are not directly comparable with the small dots. c, Enlargement of the low-identifiability region with the equivalence bounds of ±3 points. Error bars are 95% confidence intervals from 2,000 crossed-bootstrap draws for the development datasets and 10,000 for the prospective datasets; ds000117 shows the 90% intervals of its equivalence test.

Finally, we asked whether the evaluation target changes the models that researchers would select. In a nested design on FACED, we chose among nine fixed configurations using either an inner shared-material or an inner new-material validation, and evaluated both choices on the same outer new-material test. Shared-material validation selected CBraMod in all 45 outer folds, whereas new-material validation selected differential-entropy features in 36 of them (Fig. 6d). Because the candidates differed little on new materials on this task, both choices reached similar outer accuracy (Fig. 6e). The evaluation target reliably redirects model choice. What that costs grows with how far the candidates differ on new materials.

**Fig. 6.**
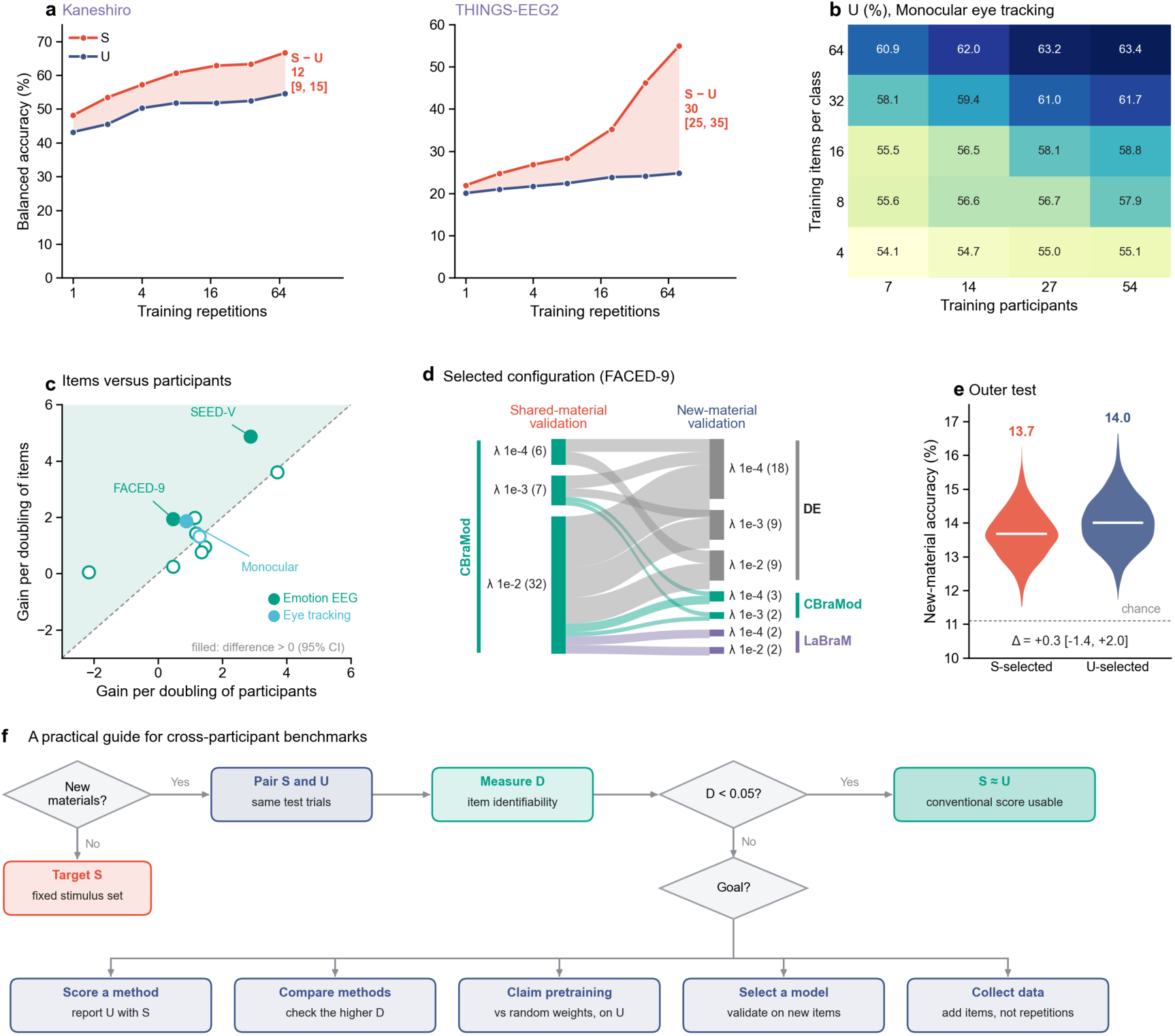
Consequences for data collection and model selection. a, Balanced accuracy under S (red) and U (blue) against the number of training repetitions in THINGS-EEG2 and Kaneshiro; labels give S − U and its 95% confidence interval at the largest number of repetitions. b, Accuracy under U in the Monocular eye-tracking study for each combination of training items per class and training participants. c, Gain in U per doubling of training items per class against the gain per doubling of training participants in eleven sources, estimated with equation (4); the dashed diagonal marks equal gains, and filled symbols mark sources in which items gained more, with a 95% interval for the difference above zero. d, Configurations selected by inner shared-material and inner new-material validation in 45 outer folds on FACED-9; band widths give the number of folds, and λ is the L2 penalty. e, Outer new-material accuracy of the configurations chosen by each validation; Δ is their difference with its 95% confidence interval, and the dashed line marks chance. f, Practical guide that follows from these results. The boxes summarize the paired evaluation (Fig. 1a), the measurement of item identifiability D (Fig. 4a), the agreement of S and U where D was below 0.05 (Fig. 5a,c) and the evidence for each goal: scoring a method (Fig. 3a), comparing methods (Fig. 4c), claiming a pretraining gain (Fig. 3c,d), selecting a model (Fig. 6d) and collecting data (Fig. 6a,c). Intervals come from 2,000 crossed-bootstrap draws over participants and items.

## Discussion

Cross-participant evaluation is the main test of whether a neural decoder generalizes to new people. When participants share experimental materials, this test also rewards recognition of the materials, by an amount set by how well individual items can be identified across people. For the film clips of FACED, where identifiability was high, shared materials added more than 20 points of accuracy and reversed the advantage of foundation models over handcrafted features. For briefly presented faces, where it was near zero, both evaluations gave the same verdict. The long, vivid materials that make a dataset attractive for cross-participant decoding also make its shared-material benchmark most misleading.

Naturalistic materials evoke responses that are shared across people. Viewers of the same film show synchronized activity across much of the cortex^35^, functional alignment builds a common space from responses to the same materials^5,36^, contrastive methods align EEG across people^24^, and individual natural images can be identified from brain activity^18^. A decoder trained on many participants can thus learn item responses along with class structure, and exploit them in a held-out participant who experienced the same items. Representations that keep more item-specific detail gain more, so a more expressive model can look better on a shared-material benchmark without being better on new materials. When materials are brief, repeated only once or twice or similar within a class, little item-specific signal survives and the two evaluations agree, as in ERP CORE and ds000117. Because the argument concerns what a representation retains, it applies to any decoder trained on shared materials, including the models of eye movements in our data.

Eye movements, which are also reproducible across viewers, may carry part of the item-specific signal in EEG, consistent with the peak of identifiability over frontotemporal sites. They do not explain the bias away. Posterior channels retained most of the identifiability, fMRI responses without ocular signal also showed a shared-material gain, and a decoder that recognized items from eye movements would profit from shared materials all the same.

Psychology recognized half a century ago that conclusions from a fixed set of materials need not extend to new ones and that items, like participants, should be treated as a random factor^15–17^. Visual neuroscience asked decoders to identify images whose responses had never been measured^18^. Neural decoding has adopted the participant half of this lesson far more often than the item half. Our crossed design and crossed bootstrap supply the item half, much as genomics keeps groups of dependent examples wholly in training or test data^20^. Earlier EEG studies showed that shared materials raise accuracy^22,23^. We show that the inflation differs across methods, can reverse comparisons and can be anticipated from item identifiability. It resembles the data leakage that inflates predictive performance in neuroimaging and genomics^19,20^, but it arises from experimental design and so persists in carefully constructed benchmarks. Concerns that benchmark gains of foundation models reflect familiarity with the evaluation data^21^ or shortcuts that do not transfer^37^ take a measurable form here, much as relating model outputs to evolutionary information explained why larger protein language models do not always predict fitness better^38^.

These findings suggest three changes to current practice (Fig. 6f). First, cross-participant benchmarks should report accuracy on new materials alongside the conventional score, as recent foundation models of neural activity have done by testing on new stimulus domains^39^. Shared-material accuracy remains the right target only when a decoder will meet just the materials it was trained on, as with the fixed stimulus set of a P300 speller. In our data, the shared-material benchmark overstated what EEG foundation models add over handcrafted features, whereas new-material accuracy revealed a smaller but real benefit of pretraining. Second, item identifiability, measurable before any method is compared, indicates whether the conventional score can be trusted. Identifiability below about 0.05 accompanied shared-material gains within a few points, and gains of 20 points or more occurred only where it exceeded 0.3.

Third, decoders meant to generalize to new materials need more items, not only more participants. Doubling the items raised new-material accuracy more than doubling the participants in three of eleven sources and never clearly less. This agrees with evidence from protein engineering that data diversity, beyond data size, governs how well models generalize^40^, and with recent neural datasets that sample thousands of images per participant^41^. Large samples of participants remain necessary for studies of individual differences^42^, but they do not substitute for samples of materials.

Overall, our work explains why cross-participant benchmarks can rank decoders differently from how they perform on new materials, and when they do not. The paired evaluation applies unchanged to any decoder. Prospective tests on highly identifiable datasets will show how precisely identifiability predicts the size of the bias, and larger models trained end to end on more diverse materials will show how far pretraining at scale closes the gap. The issue is likely to extend to any model trained on human responses to shared materials, from speech decoding to models of behaviour and annotation.

## Methods

### Data sources and scope

We focused on data sources that allow the contribution of shared materials to be isolated. A source was included if it met three conditions. First, every participant experienced the same experimental materials, so that a held-out participant is tested on items the model may have seen in other people. Second, the class label of a trial was determined by the item presented, so that withholding items does not change the label distribution. Third, each class contained several items, so that items could be partitioned into folds within each class. Sources in which labels came from each participant’s own report, or in which each item was shown to a single participant, cannot separate transfer to new people from transfer to new materials; we did not use them for the main analyses. All datasets were recorded by their original investigators with informed consent and ethical approval. We analysed their de-identified public releases and collected no new data.

Emotion EEG. We used seven public datasets in which participants watched film clips or music-video excerpts selected to elicit target emotions: FACED^26^, SEED^6^, SEED-IV^43^, SEED-V^44^, SEED-VII^45^, DEAP^25^ and DREAMER^46^. The label of a trial is the designated category of its clip, as in the published cross-participant benchmarks of these datasets. This gave ten tasks, named by dataset and number of classes: the nine designated categories of FACED (FACED-9) and its valence contrast (FACED-2), the categories of SEED, SEED-IV, SEED-V and SEED-VII, the valence–arousal quadrant of each DEAP video (DEAP-4) and its valence (DEAP-2), and the nine target emotions of DREAMER (DREAMER-9) and their valence (DREAMER-2) (Supplementary Table 1).

Visual EEG. THINGS-EEG1 presented 200 validation images 12 times to each participant in rapid serial visual presentation^27^; we used 48 participants and the 57 images belonging to the five high-level THINGS categories with at least eight images. THINGS-EEG2 presented its test images 80 times to each of ten participants^31^; 88 images in nine categories met the same criterion. The Kaneshiro dataset presented 72 images from six categories 72 times to each of ten participants^28^. Epochs ran from −0.2 to 0.8 s around image onset, were down-sampled to 100 Hz and baseline-corrected on the pre-stimulus interval, using the preprocessed data released by the authors.

fMRI. The COCO part of the Natural Object Dataset presented the same 120 natural images to nine participants, once per run^29^. We used runs 1–10, giving ten repetitions per image, and averaged the released single-trial response estimates within each of the 400 cortical parcels of the Schaefer atlas^47^. The label of an image is the COCO supercategory with the largest summed annotation area; 90 images in four supercategories with at least eight images each were retained.

Eye tracking. We used four free-viewing studies from a public eye-movement database^30^: Baseline (48 participants), Memory I (35), Tactile (57) and Monocular (68). Images belonged to two to five study-defined categories. We kept only images seen by every participant of a study, so that participants and items were fully crossed. Each viewing was represented by the duration-weighted distribution of fixations on a 16 × 12 grid, excluding the first fixation.

Prospective datasets. Three EEG datasets entered only the prospective tests described below. ERP CORE N170 has 40 participants and 160 images of faces, cars and their scrambled versions, each presented twice^32^. In EmoEEG-MC, 56 participants completed all 21 video trials of seven emotion categories^33^. In the face-recognition dataset ds000117, 16 participants saw 240 images of famous, unfamiliar and scrambled faces twice each^34^.

### Paired evaluation of shared and new materials

Within each class, items were partitioned into F folds, and participants into five folds (four for ds000117; Supplementary Table 1). A test cell (s, f) tests the participants of fold s on the items of fold f. The new-material arm U trains on the remaining participants’ trials of items outside fold f, so that every test item is new to the model. Each shared-material arm S_g_ (g ≠ f) trains on the same participants’ trials of items outside fold g. It therefore contains the test items, with the number of training items per class differing from U by at most one. All arms are scored on identical test trials, and the shared-material gain is

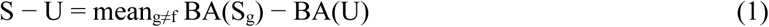

where BA is balanced accuracy. A conventional arm, Full, trains on all items of the training participants, as published benchmarks do. We used it to rank representations as a conventional benchmark would and to compare these rankings with those under U. The number of item folds F for each source is given in Supplementary Table 1. Each arm uses the same training participants and the same repetitions or windows per item. Items were dealt to folds in turn within each class, so the arms differ by at most one training item per class, and by none in the strictly budget-matched design. A trial is one participant experiencing one item; its prediction averages the log-probabilities of its windows or repetitions before the class with the largest mean is chosen. Three seeds were used throughout, and each seed draws new participant and item folds. Each pair of S and U scored on the same test trials for one dataset, representation and analysis setting counts as one matched comparison; the analyses reported here, including controls and prospective tests, comprise 318 such comparisons. For FACED, the comparison in Fig. 3a used a strictly budget-matched design. Every arm trained on two clips per class with the same participants and number of windows, and all 28 test clips and 123 participants were kept.

### Representations and readouts

Differential entropy (DE) in five bands (1–4, 4–8, 8–14, 14–31 and 31–50 Hz) was computed per channel from the same non-overlapping 10-s windows that the foundation models received (same-window DE). Official DE denotes the DE features released with FACED and the SEED datasets, which differ in window length and smoothing; for DEAP and DREAMER we computed DE over 1-s windows of the released recordings. Bipolar DE is same-window DE computed on the 16 bipolar derivations that BIOT receives, with or without BIOT’s division of each derivation by the 95th percentile of its absolute amplitude in each window (norm.). For visual and face EEG, the ERP representation is the epoch itself, channels by time samples.

CBraMod^9^, LaBraM^8^ and BIOT^10^ were used with their released pretrained weights and unchanged architectures. The pretraining corpus of LaBraM includes SEED and SEED-IV, whereas that of CBraMod contains none of the datasets analysed here. EEG was resampled to 200 Hz and expressed in units of 100 µV. Frozen representations of CBraMod and LaBraM average the last-layer patch embeddings over patches for each channel, giving C × 200 dimensions for C channels; their pooled variants also average over channels. BIOT receives the bipolar derivations as 1-s short-time Fourier windows and returns a 256-dimensional representation. Single epochs of visual and face EEG were resampled to 200 Hz and treated as one patch per channel. Together with one randomly initialized CBraMod, these choices gave the ten representations compared across the emotion tasks.

Unless stated otherwise, representations were read out by multinomial logistic regression with an L2 penalty of λ = 10^−3^, full-batch L-BFGS with up to 300 iterations and a class-balanced loss. On THINGS-EEG1 and in fMRI, this logistic readout was compared with a nearest-centroid readout, which assigns a trial to the class whose mean training response is closest in Euclidean distance after the same standardization. Eye-tracking fixation maps and fMRI parcel responses were read out with the same logistic regression. As in the published benchmarks, features were z-scored within each participant and session over all of that participant’s windows, without labels, and this applied to every arm, including the strictly budget-matched comparison. Nested model selection, the directional identifiability test, the channel analysis and the prospective tests instead took normalization statistics from training data only. The reversal on FACED-9 persisted when the statistics left out the test items (Extended Data Fig. 5a).

Pretraining controls. Randomly initialized CBraMod and LaBraM networks were embedded and read out exactly as the pretrained networks. One random CBraMod entered the ten-representation comparison. Three random networks per architecture, drawn with initialization seeds 0, 1 and 2, gave 60 frozen comparisons with the pretrained networks over the ten emotion tasks. Fine-tuned comparisons used linear probing followed by full fine-tuning (LP-FT)^12^. A two-layer head was first trained on the frozen backbone for five epochs; the whole network was then trained for five epochs with learning rates of 10^−5^ for the backbone and 10^−4^ for the head. On FACED-9, FACED-2 and SEED-V, hyper-parameters were also selected by nested validation inside each test cell, without access to its test participants or test items. Randomly initialized networks were tuned on the same inner splits (Supplementary Note 1).

Nested model selection. On FACED, nine configurations (same-window DE, CBraMod and LaBraM, each with λ of 10^−4^, 10^−3^ or 10^−2^) were compared inside each of the 45 outer training pools of the strictly budget-matched design. The outer training participants were split into two halves and the two training clips per class into two groups, and every configuration was fitted on each combination. Each model was scored on the held-out half both on the clips it was trained on (inner shared-material validation) and on the other clips (inner new-material validation). One configuration was selected by each validation, refitted on the whole outer training pool and scored on the common outer new-material test.

### Control analyses

Further controls in emotion EEG used official DE and the logistic readout unless stated otherwise. The first asked whether new-material accuracy profited from clips cut from the same source film as a test clip. An arm U_film_ removed from the training pool of U every clip that shared a source film with a test clip, and an arm U_rand_ removed the same number of clips per class at random. On FACED-9, SEED and SEED-IV, only classes whose clips came from more than one film were used, and this control used one seed. SEED-IV, SEED-V and SEED-VII presented their clips in a fixed order. For official DE and frozen CBraMod, we compared new-material accuracy on test trials whose presentation position matched that of training clips of the same class with accuracy on trials whose position did not. To test whether the gain depended on how strongly a clip was felt, we ranked the viewers of each clip in FACED-9, SEED-V and SEED-VII by their own ratings of it. The accuracy of the upper third was compared with that of the lower third, under U and under S. On DEAP and DREAMER, the designated valence of each clip was replaced by each participant’s own valence rating, split at the midpoint of the rating scale, for each of eight representations.

The remaining controls changed the protocol. Held-out normalization computed the statistics for each participant and session only from the windows of clips outside the test-item fold. A multilayer perceptron replaced the logistic readout. It had hidden layers of 256 and 128 units, layer normalization, GELU activations, dropout of 0.5 and input dropout of 0.2, and was trained with AdamW (learning rate 10^−3^, weight decay 10^−2^, one-cycle schedule) for 3,000 steps with batches of 1,024 windows and a class-weighted cross-entropy loss. Finally, the L2 penalty of the logistic readout was varied from λ = 10^−4^ to 1. Intervals for all controls come from 2,000 crossed-bootstrap draws.

Method comparisons. For a representation A and a comparator B evaluated on identical test trials,

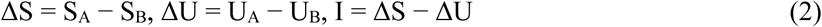

A positive interaction means that the advantage of A is larger on shared materials than on new materials, and opposite signs of ΔS and ΔU indicate a reversal. Comparisons with handcrafted features estimate the combined difference in representation, architecture and learning, whereas comparisons between pretrained and randomly initialized networks of the same architecture estimate the effect of pretrained weights^48^. To compare rankings, the ten representations were ranked within each emotion task by their accuracy under Full and under U.

### Item identifiability

Within each class, the logistic readout described above was trained to identify the class’s items from the trials of one set of participants and tested on held-out participants, averaging the log-probabilities of a trial’s windows or repetitions. With n items in a class and trial accuracy A, the chance-corrected identifiability is

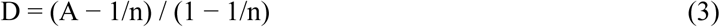

which is 0 at chance and 1 for perfect identification. D was averaged within classes and then across classes. For the correlation with the shared-material gain across the 100 representation–task pairs of emotion EEG, identification used five-fold participant cross-validation over all items. For the directional test on six representation pairs, probes were fitted inside every new-material training pool only. Its training participants were split into two halves that alternated as training and validation sets, only the items available to U were used, and the normalization statistics came from the training half.

Removal of item-identity directions. For each test cell, the differences between the training item means and the mean of the item means of their class were estimated from the training windows of U, which contain no test participant and no test item. Their orthonormal basis, obtained by singular value decomposition, was projected out of every window, and the readouts were retrained and scored as in the main analysis. A control removed the same number of random orthonormal directions drawn independently for each test cell.

Channel analysis. We recomputed D for same-window DE on FACED from subsets of channels. Besides all 32 channels, we used the 24 channels remaining after removing the frontopolar, lateral frontal, temporal and mastoid channels (Fp1, Fp2, F7, F8, T7, T8, A1 and A2), 14 centro-parietal, parietal and occipital channels, ten random subsets of 14 channels and each of the 32 channels alone, the last using the five DE bands of one electrode. According to the dataset documentation, the two channels labelled HEOL and HEOR in the second FACED cohort were placed on the mastoids and correspond to A1 and A2 of the first cohort. Participants were split into halves five times, each half serving once for training, with normalization statistics from the training half.

### Prospective tests

Each prospective test followed the same sequence. The data definition was fixed, and its document and code were hashed and time-stamped before any recording was read. The forecasting rule, the probe and the decision criteria were then fixed and hashed. The identifiability probes were run on new-material training pools only, and the resulting forecast was written to a hashed file. Category decoders were trained only after this file existed, and the analysis code refused to fit them otherwise.

The directional rule forecast sign(I) = sign(D_A_ − D_B_) when the 95% interval of D_A_ − D_B_ excluded zero and abstained otherwise. For ds000117, the rule was extended before any probe was run. If the upper 95% limits of D for both representations were below 0.05, it issued a null forecast: the shared-material gain of each representation and their interaction would lie within ±3 points. The null forecast was judged supported if the 90% intervals of all three quantities lay within (−3, 3) points, an equivalence test at a one-sided α of 0.05^49^. It was judged contradicted if any 95% interval excluded zero with an estimate beyond ±3 points.

ERP CORE epochs were taken from −0.2 to 0.8 s around image onset after average referencing and 0.1– 30-Hz band-pass filtering, and down-sampled to 100 Hz. Intact and scrambled versions of the same image were kept in the same item fold. EmoEEG-MC used the released preprocessed recordings (0.1–47 Hz, independent component analysis, average reference, 200 Hz), whose 30-s trials were cut into three 10-s windows as for FACED. ds000117 used the EEG channels of the released recordings with average referencing, 0.1–30-Hz band-pass filtering and epochs from −0.2 to 0.8 s at 100 Hz, keeping the 80 images per class that every participant saw exactly twice. The representation pairs were CBraMod versus ERP for ERP CORE and ds000117, and CBraMod versus same-window DE for EmoEEG-MC.

### Training repetitions and the scaling of items and participants

To test the effect of repeated presentations, we compared new-material and shared-material accuracy when training on all repetitions with training on a random subset of repetitions, keeping the test repetitions unchanged. To compare items with participants as sources of training data, we kept each test fold fixed, trained the new-material arm on m items per class from N participants over a grid of m and N, and fitted

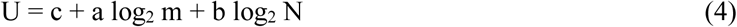

so that a and b are the gains per doubling of items and of participants. In nine emotion tasks, m ranged from one to at most 16 training items per class and N from three to 98 training participants, depending on the dataset. Each grid point used two random draws; DREAMER-9 has only one training item per class and was not used. In the two largest eye-tracking studies, m took the values 4, 8, 16 and 24 items per class in Tactile and doubled from 4 to 64 in Monocular. N took four roughly doubling values up to all training participants, with three seeds. In each bootstrap draw, the accuracy of every grid cell was recomputed with the resampled participants and items before equation (4) was refitted, and the difference a − b was taken from the same draw.

### Statistics

Participants and items are both random factors in these designs. Intervals were therefore obtained with a crossed bootstrap that resamples participants and, within each class, items with replacement, applying the same draw to all methods and arms being compared. We used 2,000 draws for the development analyses and 10,000 for the prospective tests. In simulations that retained the participant, class and item counts of each emotion task, the nominal 95% intervals covered the true gain in 89.9% to 99.9% of simulated datasets (Supplementary Note 2 and Supplementary Table 4). Intervals are percentile intervals conditional on the fitted models. Correctness was pooled per participant and item over seeds and arms before balanced accuracy was computed, so that seeds and arms were not treated as independent observations. Correlations between D and the shared-material gain are Spearman coefficients over representation–task pairs. Pairs from one task share data, so the coefficients are descriptive; within-task versions subtract each task’s mean before correlating. The authors implemented all analyses in Python 3.11.16 with NumPy 2.4.6, SciPy 1.17.1, PyTorch 2.12.1 and MNE-Python 1.13.2, and a large language model from Anthropic assisted in revising and refining the code.

## Data availability

This study analysed only previously published datasets, which are available from their original repositories: FACED^26^ (Synapse, syn50614194), SEED^6^, SEED-IV^43^, SEED-V^44^ and SEED-VII^45^ (https://bcmi.sjtu.edu.cn/home/seed/, available on request to the dataset owners), DEAP^25^ (https://www.eecs.qmul.ac.uk/mmv/datasets/deap/, on request), DREAMER^46^ (Zenodo, https://doi.org/10.5281/zenodo.546113, on request), THINGS-EEG1^27^ (OpenNeuro, ds003825), THINGS-EEG2^31^ (OSF, https://osf.io/3jk45/), the Kaneshiro dataset^28^ (Stanford Digital Repository, https://purl.stanford.edu/bq914sc3730), the Natural Object Dataset^29^ (OpenNeuro, ds004496), the free-viewing eye-movement database^30^ (Dryad, https://doi.org/10.5061/dryad.9pf75), ERP CORE^32^ (OSF, https://osf.io/thsqg/), EmoEEG-MC^33^ (OpenNeuro, ds005540) and the face-recognition dataset ds000117^34^ (OpenNeuro, ds000117). No new data were collected for this study. Source data are provided with this paper.

## Code availability

The custom code used for all analyses and figures is available from the corresponding author upon request.

## Author contributions

H.C. conceived the study, designed the paired evaluation protocol, implemented the analysis pipeline, performed the experiments and analyses, and wrote the manuscript. Y.Z. and S.H. contributed to the analysis and interpretation of the results and revised the manuscript. T.X. supervised the project, guided the study design and revised the manuscript. All authors read and approved the final manuscript.

## Extended Data

**Extended Data Fig. 1.**
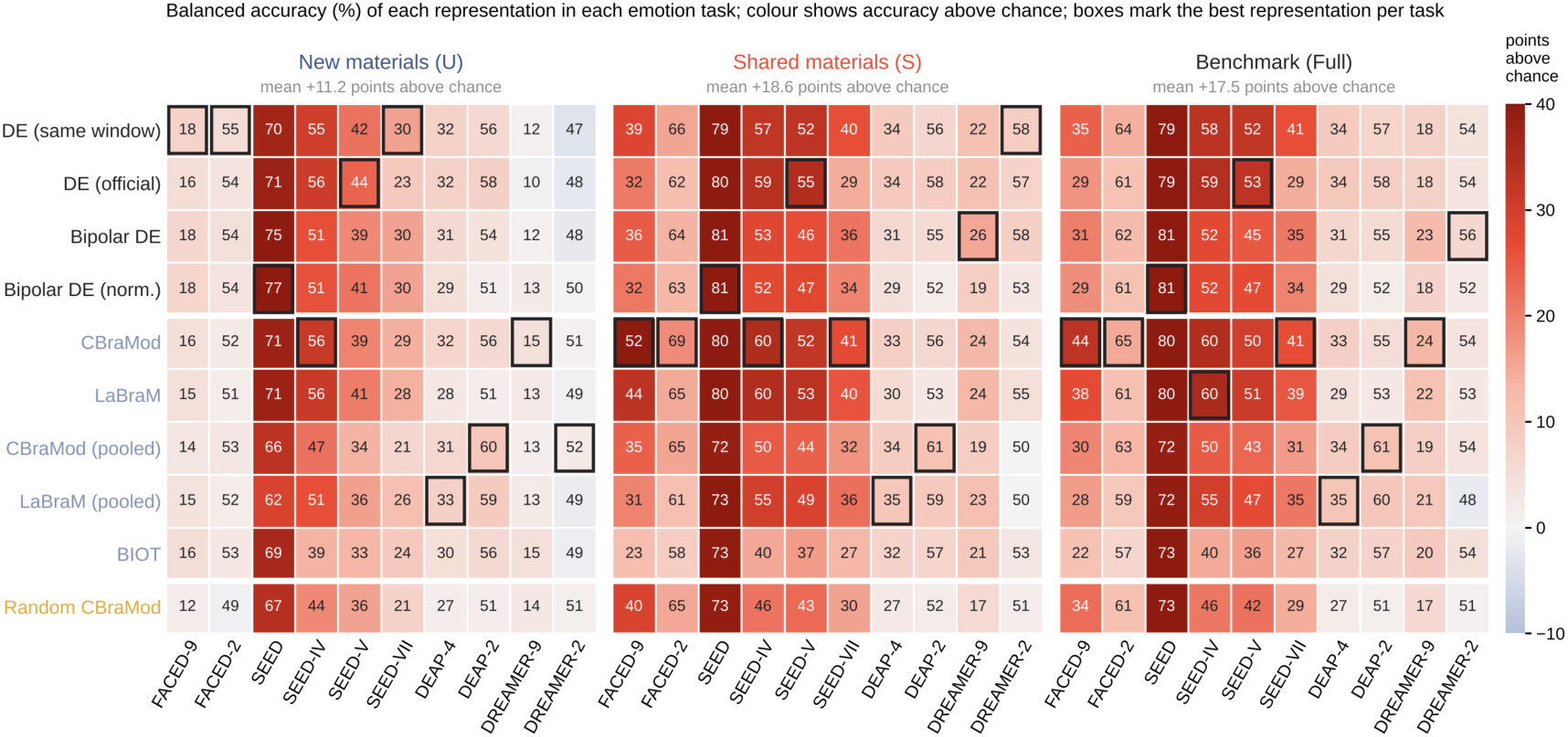
Balanced accuracy of ten representations in ten emotion tasks. Balanced accuracy (%) under new-material evaluation (U), shared-material evaluation (S) and the conventional benchmark that trains on all items (Full), for each representation (rows) and task (columns). Colour gives accuracy above chance; boxes mark the representation with the highest accuracy in each task. Values above each matrix give the mean accuracy above chance. Frozen representations with logistic readouts; three seeds.

**Extended Data Fig. 2.**
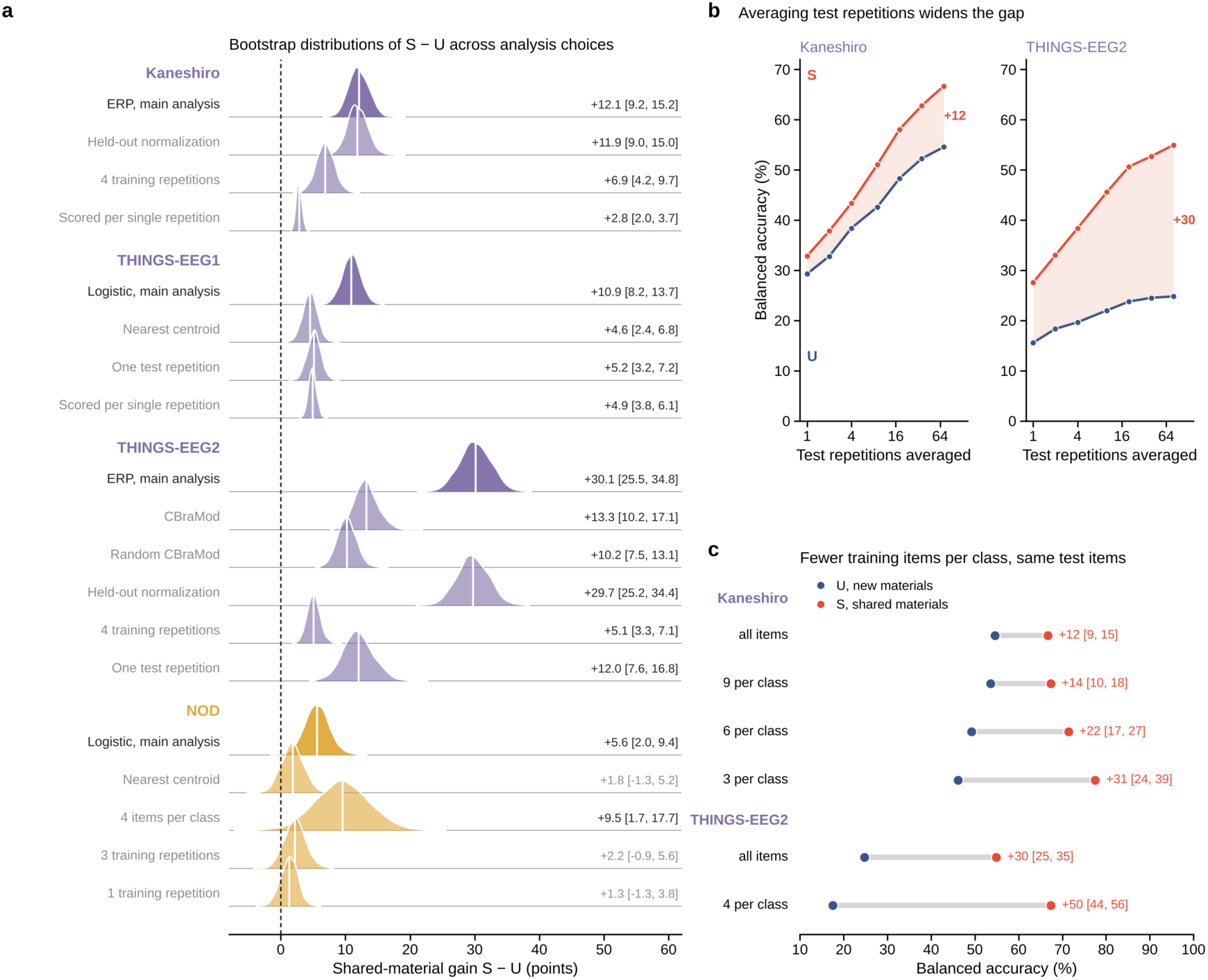
Shared-material gain in visual EEG and fMRI under alternative analysis choices. a, Bootstrap distributions of S − U for the main analysis of each source (dark) and for alternative choices of representation, readout, normalization, number of training repetitions, number of training items and scoring (light); white lines mark means, and labels give the mean and 95% interval. b, Balanced accuracy under S (red) and U (blue) against the number of test repetitions averaged, after training on all repetitions; labels give S − U at the largest number. c, Balanced accuracy under U and S after training on fewer items per class while keeping the test items fixed; labels give S − U with its 95% interval. Distributions and intervals come from 2,000 crossed-bootstrap draws.

**Extended Data Fig. 3.**
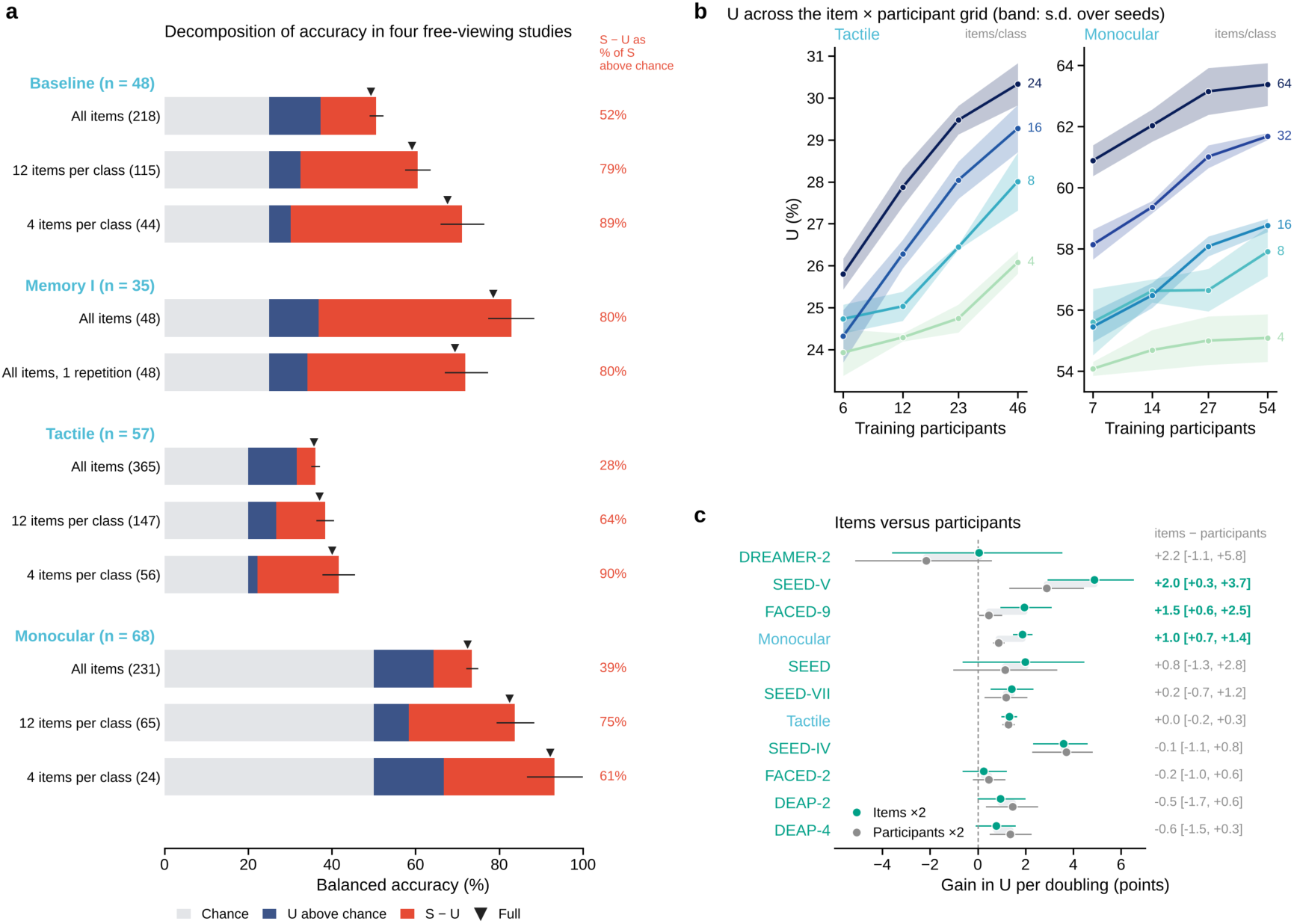
Eye tracking and the scaling of items and participants. a, Balanced accuracy in four free-viewing studies decomposed into chance (grey), accuracy of U above chance (blue) and the shared-material gain S − U (red), after training on all items, on 12 or 4 items per class, or on all items with one repetition; numbers in parentheses give the items entering the test, whiskers the 95% interval of S − U, triangles the conventional benchmark (Full), and percentages the share of above-chance S that comes from S − U. b, U across the grid of training items per class and training participants in the Tactile and Monocular studies; bands give the standard deviation over seeds. c, Gain in U per doubling of training items (teal) and of training participants (grey) from equation (4) in eleven sources, with 95% intervals; values on the right give the difference with its 95% interval, bold where it excludes zero.

**Extended Data Fig. 4.**
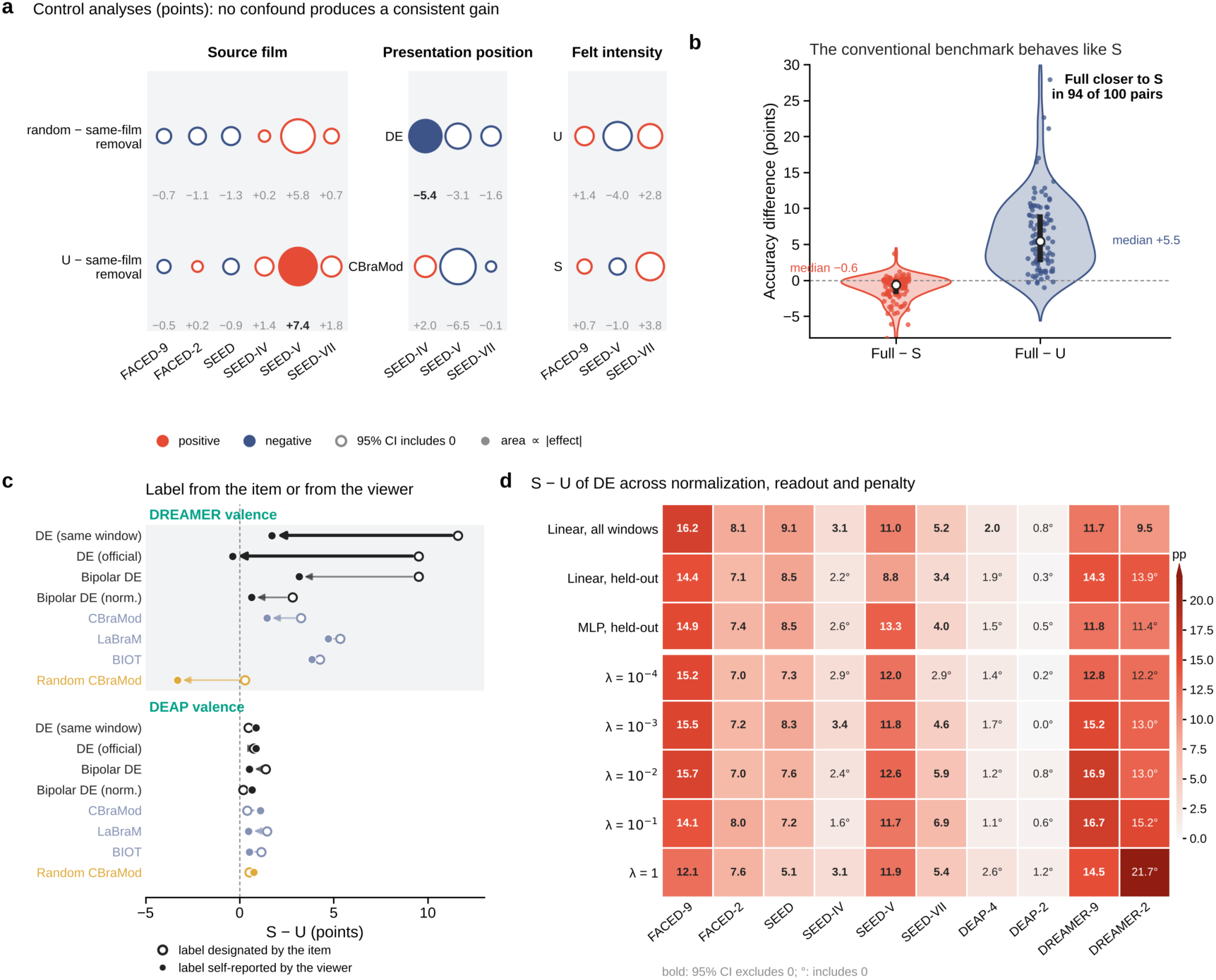
Alternative explanations of the shared-material gain in emotion EEG. a, Effects (points) of three possible confounds. Source film: U after removing, from training, the clips cut from the same source film as any test clip (same-film removal), compared with removing as many clips per class at random (top) and with the unrestricted U (bottom), with FACED-9, SEED and SEED-IV restricted to classes whose clips came from more than one film; official DE, linear readout, one seed. Presentation position: difference in U between test clips presented at the same position as their training counterparts and at a different position, in the three SEED datasets with a fixed presentation order, for DE and frozen CBraMod (three seeds). Felt intensity: difference in U and in S between the upper and lower thirds of viewers ranked by their rating within each clip. Bubble area is proportional to the magnitude of the effect, red bubbles are positive and blue negative, and filled bubbles mark 95% intervals that exclude zero; values below the bubbles give the effect. b, Difference between the conventional benchmark (Full) and S, and between Full and U, for the 100 representation–task pairs of frozen readouts; violins give the density, points the pairs and white circles the medians. c, S − U with labels designated by the item (open circles) and with labels self-reported by each viewer (filled circles) for valence in DREAMER and DEAP; arrows run from designated to self-reported labels and are thick where the 95% interval of the change excludes zero. Three seeds. d, S − U of DE in each task with a linear readout on all windows (main analysis, three seeds), with normalization statistics computed only from clips outside the test fold for linear and multilayer-perceptron readouts, and with linear readouts at five L2 penalties λ (one seed). Bold values have 95% intervals that exclude zero; ° marks intervals that include zero. Intervals are 95% intervals from 2,000 crossed-bootstrap draws.

**Extended Data Fig. 5.**
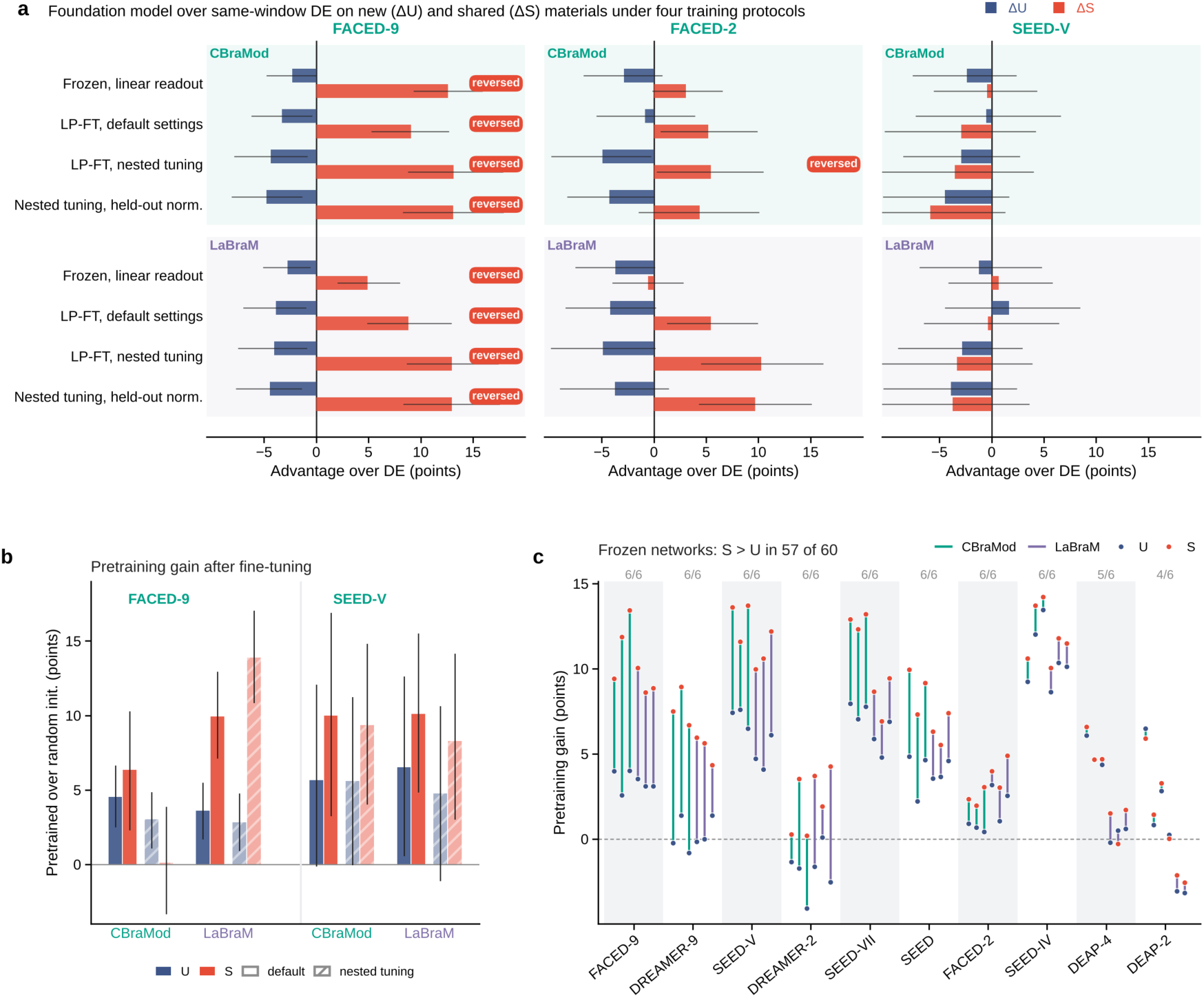
Fine-tuning, nested tuning and pretraining controls. a, Advantage of CBraMod and LaBraM over same-window DE on new materials (ΔU, blue) and on shared materials (ΔS, red) on FACED-9, FACED-2 and SEED-V, for frozen linear readouts, LP-FT with default settings, LP-FT with nested hyper-parameter tuning and nested tuning with held-out normalization; “reversed” marks comparisons in which the 95% interval of ΔS lies above zero and that of ΔU below zero. b, Advantage of pretrained over randomly initialized networks of the same architecture under U (blue) and S (red), after LP-FT with default settings (solid) and with nested tuning (hatched). c, Pretraining gain of frozen networks under U (blue) and S (red) for three random initializations of each architecture in each task (60 comparisons); numbers above each task count comparisons with a larger gain under S. Bars and points give estimates and lines 95% intervals from 2,000 crossed-bootstrap draws.

**Extended Data Fig. 6.**
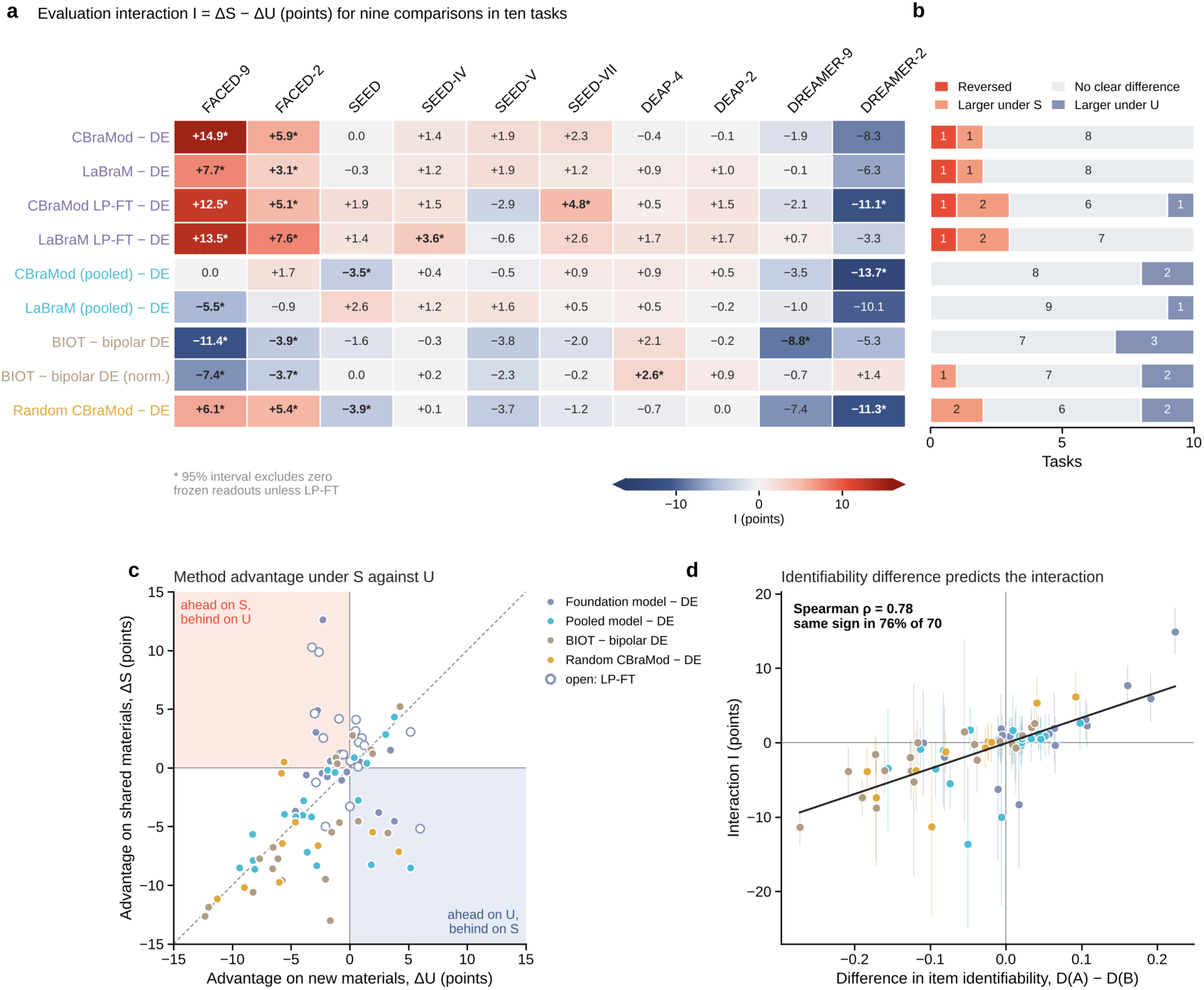
Evaluation interactions across method comparisons and tasks. a, Evaluation interaction I = ΔS − ΔU (points) for nine comparisons (rows) in ten emotion tasks (columns); asterisks mark 95% intervals that exclude zero. Readouts are frozen unless marked LP-FT. b, Number of tasks in which each comparison reversed (advantage of opposite sign under S and U, both intervals excluding zero), was larger under S, showed no clear difference, or was larger under U. c, Advantage under S against advantage under U for all 90 comparison–task pairs; the shaded quadrants contain comparisons that lead under one evaluation and trail under the other, and the dashed line marks equal advantages. d, Difference in item identifiability between the two representations against I for the 70 frozen comparison–task pairs, with a least-squares line; ρ, Spearman correlation. Error bars are 95% intervals from 2,000 crossed-bootstrap draws.

**Extended Data Fig. 7.**
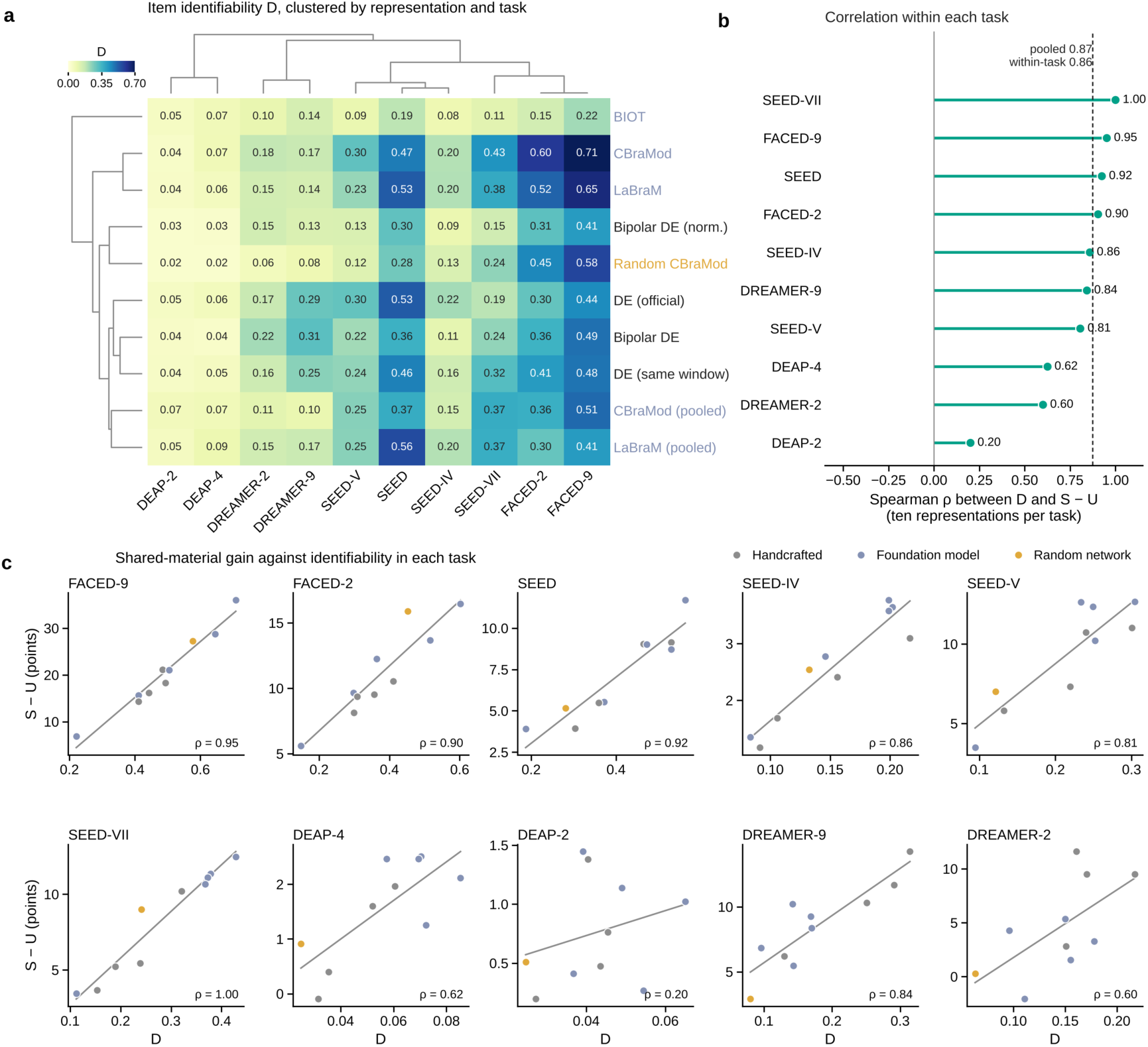
Item identifiability across representations and tasks. a, Item identifiability D for ten representations in ten emotion tasks, with representations and tasks ordered by hierarchical clustering (average linkage; Euclidean distance for representations, correlation distance for tasks). b, Spearman correlation between D and the shared-material gain S − U across the ten representations within each task; the dashed line marks the pooled correlation over all 100 pairs. c, S − U against D in each task, coloured by representation type, with least-squares lines.

**Extended Data Fig. 8.**
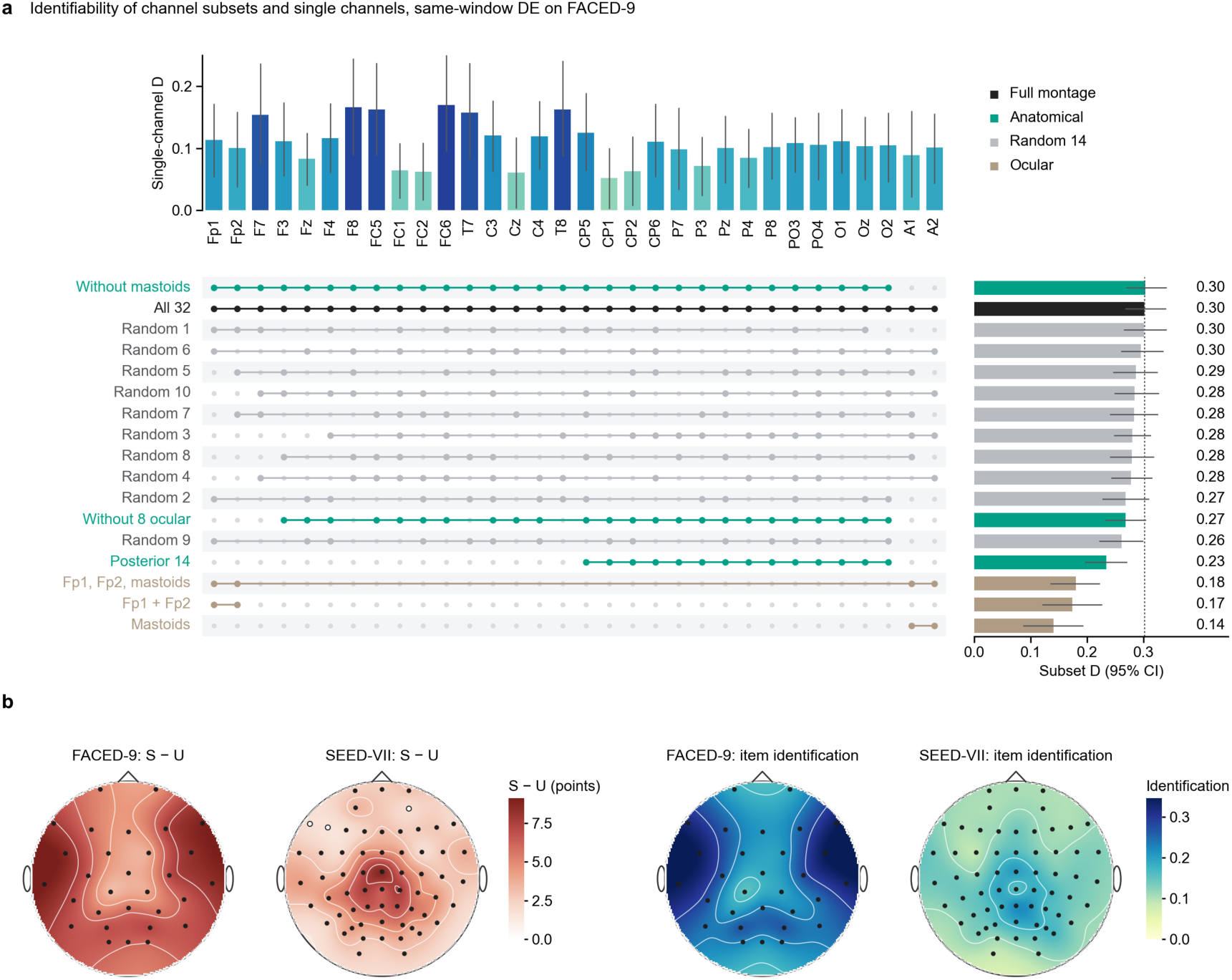
Where on the scalp the item information sits. a, Identifiability D of channel subsets and of single channels for same-window DE on FACED-9. Bars on the right give D of each subset with its 95% interval, sorted by D, and the matrix marks the channels in each subset; bars on top give D of each channel alone. Subsets are the full montage, anatomically defined subsets, ten random 14-channel subsets and subsets restricted to the channels most exposed to ocular artefacts. The dotted line marks D of the full montage. b, Electrode maps of the shared-material gain S − U (left) and of item identification (right, chance-corrected accuracy of identifying the clip) from one same-window DE readout per electrode on FACED-9 (30 scalp electrodes) and SEED-VII (62 electrodes); one colour scale per quantity across the two tasks. Values between electrodes are interpolated with thin-plate splines; filled dots mark electrodes with q < 0.05 (Benjamini–Hochberg over the electrodes of each task) and open dots the remainder. Seeds 0–2 pooled; intervals and q values come from 2,000 crossed-bootstrap draws over participants and items.

**Extended Data Fig. 9.**
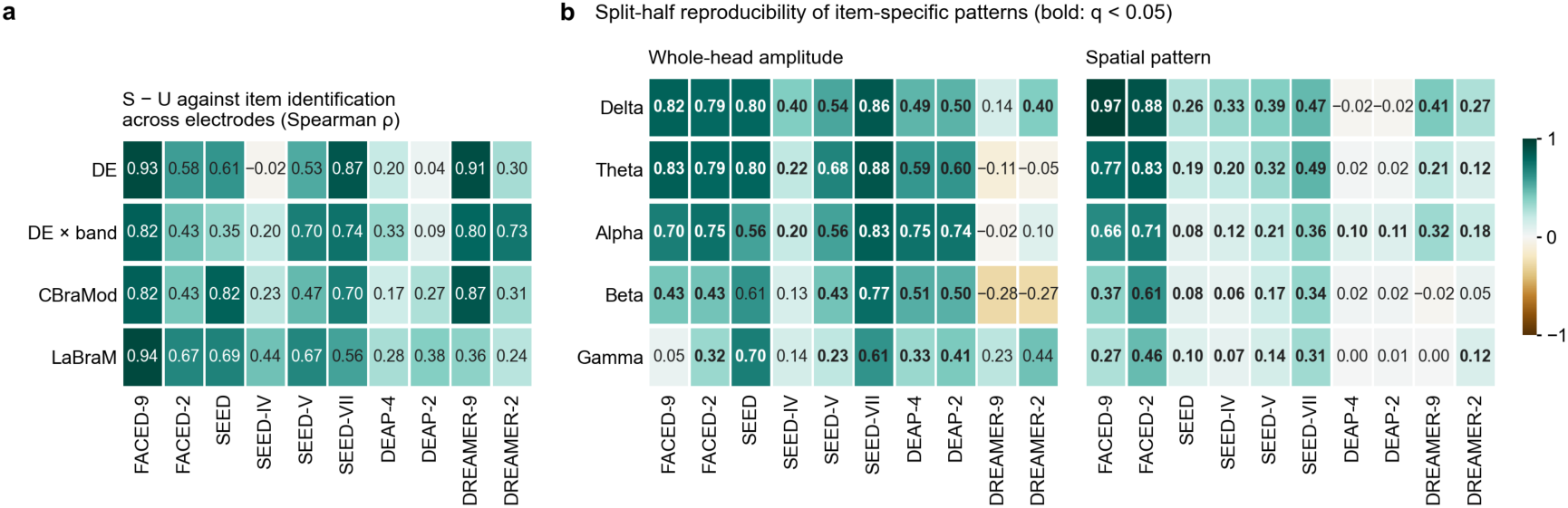
Electrode-level link and reproducibility of item patterns. a, Spearman correlation across electrodes between the per-electrode S − U and item identification in each task, for one DE readout per electrode (five bands), one readout per electrode and band, and the channel embeddings of CBraMod and LaBraM. b, Split-half reproducibility, across participants, of item-specific patterns of same-window DE in five bands: whole-head amplitude (correlation, between participant halves, of each clip’s whole-head mean deviation from its category) and spatial pattern (correlation across electrodes of the same clip minus that of different clips of the same category, whole-head mean removed). Bold values have q < 0.05 from within-category permutation tests (1,000 permutations; Benjamini–Hochberg over the 50 cells of each statistic). Seeds 0–2 pooled.

**Extended Data Fig. 10.**
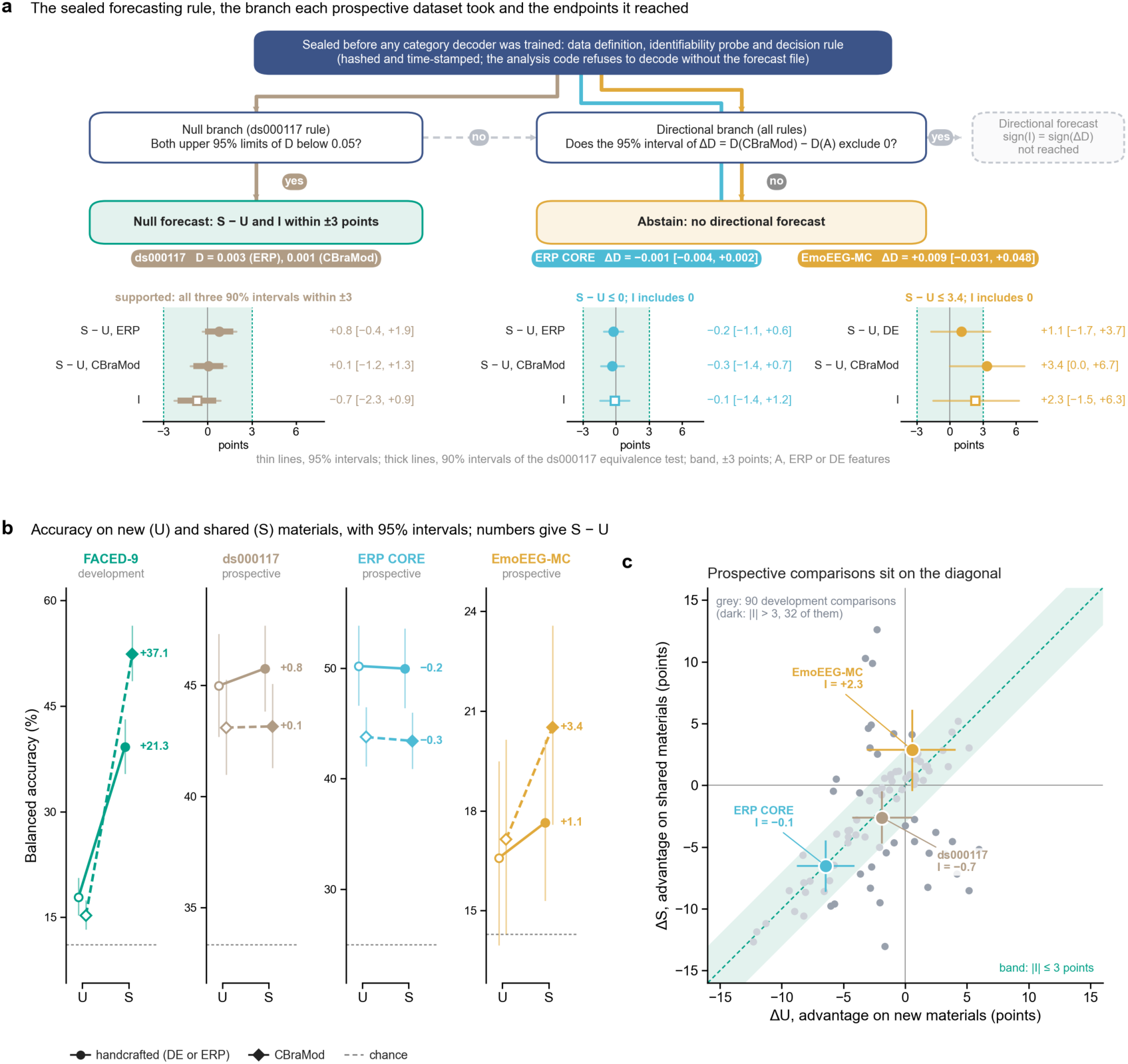
Sealed forecasting rule and prospective endpoints. a, Decision rule fixed before any category decoder was trained, with the route each prospective dataset took (coloured lines) and its sealed identifiability values. The null branch was added to the rule for ds000117 before its identifiability probe was run; ERP CORE and EmoEEG-MC were evaluated with the directional rule only, and no dataset reached a directional forecast. Under each leaf are the shared-material gain of each representation (circles) and the evaluation interaction I (squares) of the datasets that reached it, with 95% intervals (thin lines) and, for ds000117, the 90% intervals of its equivalence test (thick lines); numbers on the right give each estimate with its 95% interval, and the band marks ±3 points. b, Balanced accuracy under U (open symbols) and S (filled symbols) for the handcrafted representation (circles, solid lines; DE or ERP features) and CBraMod (diamonds, dashed lines) in the three prospective datasets and, for reference, in FACED-9 as in Fig. 5a; vertical lines give 95% intervals, dashed horizontal lines chance, and numbers S − U. c, Advantage on shared materials (ΔS) against advantage on new materials (ΔU) for CBraMod over the handcrafted representation in the three prospective datasets (coloured, with 95% intervals) and for the 90 method comparisons in emotion EEG (grey; darker where |I| > 3 points); the band marks |I| ≤ 3 points around the diagonal. Prospective intervals come from 10,000 crossed-bootstrap draws.

