## supplementary_information for "Shared experimental materials inflate and reorder cross-participant decoding benchmarks"

### Contents

Supplementary Note 1. Fine-tuning and nested hyper-parameter selection

Supplementary Note 2. Calibration of the crossed bootstrap

Supplementary Table 1. Data sources and evaluation designs

Supplementary Table 2. Robustness of the association between item identifiability and the shared-material gain

Supplementary Table 3. BIOT against bipolar differential-entropy features

Supplementary Table 4. Calibration of the crossed bootstrap

Supplementary References

S denotes shared-material evaluation, U denotes new-material evaluation, and D denotes chance-corrected item identifiability. The evaluation interaction is $I = \Delta S - \Delta U$, where $\Delta S$ and $\Delta U$ are the paired advantages of one method over its comparator.

### Supplementary Note 1. Fine-tuning and nested hyper-parameter selection

#### Default fine-tuning

Fine-tuned models used linear probing followed by full fine-tuning (LP-FT)[1]. The classification head had two layers, with dimensions C × 200 → 256 → K for CBraMod and LaBraM and 256 → 256 → K for BIOT. Here C is the number of channels and K is the number of classes. The head used ELU activation and dropout 0.1. It was trained for five epochs on the frozen backbone at a learning rate of $5 \times 10^{-4}$. The whole network was then trained for five epochs, with learning rates of $10^{-5}$ for the backbone and $10^{-4}$ for the head.

Training used AdamW with weight decay 0.05, a one-cycle schedule with 10% warm-up, label smoothing 0.1, batch size 64 and bfloat16 mixed precision. The gradient norm was clipped at 1.0 during whole-network training. Pooled embeddings were standardized within each participant-session, and the statistics were recomputed from the final network before testing.

#### Nested hyper-parameter selection

On FACED-9, FACED-2 and SEED-V, hyper-parameters were selected within each outer test cell, without using its test participants or test items. The next participant fold supplied the validation participants. For each remaining item fold, the model was trained on the other training participants' trials outside that fold and the test-item fold. It was then scored on the validation participants' trials from the held-out inner item fold. This gave two inner rotations per outer cell. Each configuration was evaluated in both rotations with two search seeds, and selection maximized mean validation balanced accuracy. Ties favoured the lower-indexed configuration, starting with the default schedule.

The initial grid for each pretrained model contained eight configurations: four learning-rate settings for whole-network training, crossed with 5 or 15 epochs. The rates spanned the default schedule and the published fine-tuning rate; the three higher LaBraM settings used layer-wise decay 0.65. If the best configuration used the longest evaluated duration and exceeded the next shorter duration at the same rates by at least one percentage point, the next duration was evaluated. This rule allowed extensions from 15 to 30 and then 50 epochs, using the same inner rotations and search seeds. Differential-entropy baselines had their L2 penalty selected on the same inner splits.

#### Random initialization and normalization controls

Randomly initialized CBraMod and LaBraM were trained on FACED-9 and SEED-V. Their default fits used the same LP-FT procedure as the pretrained networks. For nested selection, they were trained end to end from the first epoch. The grid crossed learning rates of $5 \times 10^{-5}$, $10^{-4}$, $2 \times 10^{-4}$ and $3 \times 10^{-4}$ with 15 or 30 epochs. A 50-epoch extension used the same one-point improvement rule. Inner splits, search seeds and the selection criterion matched those of the pretrained models. Participant-session normalization statistics were recomputed from the current network at the start of each epoch and before testing.

The held-out-normalization sensitivity analysis in Extended Data Fig. 5a reused the selected configurations without retuning. Within each participant-session, normalization statistics used only windows from clips outside the test-item fold. If a session contained no eligible windows, statistics were estimated from that participant's eligible windows in the other sessions.

All comparisons involving nested selection kept the participant and item folds generated by split seed 0. The three training seeds changed model initialization and training order while leaving those folds fixed. For each test cell, the shared-material arm withheld the next item fold, and each comparator was scored on the same arm and the same test trials. These comparisons are shown in Extended Data Fig. 5.

### Supplementary Note 2. Calibration of the crossed bootstrap

The crossed bootstrap resamples participants and, within each class, items. Several emotion tasks contain only two to five items per class, so we assessed its calibration with simulations that retained each task's participant, class and item counts. For differential-entropy features, we generated the participant × item matrix of differences between S and U correctness as the sum of a class effect, a participant effect, an item effect and a residual. Participant and item standard deviations were estimated by the method of moments. Residuals were resampled from the observed residuals to retain their empirical distribution.

Four scenarios crossed a class effect of zero or the observed class gain with the estimated item standard deviation or an item-heavy standard deviation of max(1.5 × estimate, 6 percentage points). Item effects were normally distributed, except in the item-heavy scenario with the observed gain, where they were resampled from the observed within-class item deviations to retain their skew. Each scenario used 2,000 simulated datasets. Each simulated dataset was analysed with 1,999 bootstrap replicates, using the same resampling procedure as the main analysis, which used 2,000 replicates.

Across the 40 task–scenario combinations, empirical coverage of the nominal 95% percentile interval ranged from 0.899 to 0.999, and absolute bias was below 0.2 percentage points. Under zero true gain, the one-sided test rejected at rates from 0.004 to 0.079. Coverage was lowest for FACED-9 and SEED in the item-heavy scenarios. The bootstrap standard error was conservative for tasks with more items per class, including SEED-IV and DEAP. Supplementary Table 4 gives the coverage and rejection rates for each task and scenario.

Supplementary Table 1. Data sources and evaluation designs

| Modality | Source or task | Participants | Classes | Item pool | Presentations per item | Item folds | Participant folds |
|---|---|---|---|---|---|---|---|
| Emotion EEG | FACED-9 | 123 | 9 | 28 | 1 | 3 | 5 |
| Emotion EEG | FACED-2 | 123 | 2 | 24 | 1 | 3 | 5 |
| Emotion EEG | SEED | 15 | 3 | 15 | 3 | 5 | 5 |
| Emotion EEG | SEED-IV | 15 | 4 | 72 | 1 | 6 | 5 |
| Emotion EEG | SEED-V | 16 | 5 | 45 | 1 | 3 | 5 |
| Emotion EEG | SEED-VII | 20 | 7 | 80 | 1 | 4 | 5 |
| Emotion EEG | DEAP-4 | 32 | 4 | 40 | 1 | 5 | 5 |
| Emotion EEG | DEAP-2 | 32 | 2 | 40 | 1 | 5 | 5 |
| Emotion EEG | DREAMER-9 | 23 | 9 | 18 | 1 | 2 | 5 |
| Emotion EEG | DREAMER-2 | 23 | 2 | 16 | 1 | 2 | 5 |
| Visual EEG | THINGS-EEG1 | 48 | 5 | 57 | 12 | 4 | 5 |
| Visual EEG | THINGS-EEG2 | 10 | 9 | 88 | 80 | 4 | 5 |
| Visual EEG | Kaneshiro | 10 | 6 | 72 | 72 | 3 | 5 |
| fMRI | Natural Object Dataset | 9 | 4 | 90 | 10 | 4 | 5 |
| Eye tracking | Baseline | 48 | 4 | 218 | 1 | 4 | 5 |
| Eye tracking | Memory I | 35 | 4 | 48 | 5 | 4 | 5 |
| Eye tracking | Tactile | 57 | 5 | 365 | 1 | 4 | 5 |
| Eye tracking | Monocular | 68 | 2 | 231 | 1 | 4 | 5 |
| Prospective EEG | ERP CORE | 40 | 4 | 160 | 2 | 4 | 5 |
| Prospective EEG | EmoEEG-MC | 56 | 7 | 21 | 1 | 3 | 5 |
| Prospective EEG | ds000117 | 16 | 3 | 240 | 2 | 5 | 4 |

The rows describe the task-specific item pools and cross-validation designs. Multiple task definitions from the same source are listed separately. Items are clips, images or scenes; presentations count repeated presentations per participant. Item folds are formed within class. The training-item and repetition subsets for individual comparisons are described in Methods and the corresponding main or Extended Data figure legends.

THINGS-EEG2 had 88 eligible images across nine categories; the evaluation selected 76 images per split seed, with 87 distinct images used across the three seeds. Thus the eligible pool and the number evaluated in one split are different counts. ERP CORE assigned intact and scrambled versions of the same source image to the same item fold. ERP CORE, EmoEEG-MC and ds000117 were used for the prospective tests.

Supplementary Table 2. Robustness of the association between item identifiability and the shared-material gain

| Analysis | Subset | Pairs | Spearman ρ |
|---|---|---|---|
| Pooled | All pairs | 100 | 0.87 |
| Pooled | Task means removed | 100 | 0.86 |
| Pooled | Ranks within each task | 100 | 0.77 |
| Single task | FACED-9 | 10 | 0.95 |
| Single task | FACED-2 | 10 | 0.90 |
| Single task | SEED | 10 | 0.92 |
| Single task | SEED-IV | 10 | 0.86 |
| Single task | SEED-V | 10 | 0.81 |
| Single task | SEED-VII | 10 | 1.00 |
| Single task | DEAP-4 | 10 | 0.62 |
| Single task | DEAP-2 | 10 | 0.20 |
| Single task | DREAMER-9 | 10 | 0.84 |
| Single task | DREAMER-2 | 10 | 0.60 |
| Single task | Median of the ten tasks | 10 | 0.85 |
| One dataset left out | Without DEAP | 80 | 0.78 |
| One dataset left out | Without DREAMER | 80 | 0.91 |
| One dataset left out | Without FACED | 80 | 0.83 |
| One dataset left out | Without SEED | 90 | 0.92 |
| One dataset left out | Without SEED-IV | 90 | 0.86 |
| One dataset left out | Without SEED-V | 90 | 0.89 |
| One dataset left out | Without SEED-VII | 90 | 0.87 |
| Difference to baseline | All pairs | 30 | 0.82 |
| Difference to baseline | Task means removed | 30 | 0.73 |
| Difference to baseline | Without FACED | 24 | 0.67 |
| Difference to baseline | Without SEED | 27 | 0.86 |
| Difference to baseline | Without SEED-IV | 27 | 0.81 |
| Difference to baseline | Without SEED-V | 27 | 0.83 |
| Difference to baseline | Without SEED-VII | 27 | 0.79 |
| Difference to baseline | Without DEAP | 24 | 0.84 |
| Difference to baseline | Without DREAMER | 24 | 0.85 |

Spearman correlations between D and S − U across representation–task pairs in emotion EEG, supporting Fig. 4b and Extended Data Fig. 7. "Task means removed" subtracts each task's mean from both quantities before correlating. "Ranks within each task" ranks the quantities separately within each task before pooling. Leave-one-dataset-out rows omit all tasks derived from the named dataset.

"Difference to baseline" correlates the difference in D with the difference in S − U for CBraMod, LaBraM and BIOT against their handcrafted comparators in ten tasks: same-window DE for CBraMod and LaBraM, and bipolar DE without normalization for BIOT. With normalized bipolar DE as the comparator of BIOT, the coefficient over all pairs is 0.68 instead of 0.82. Pairs within a task share data; these correlation coefficients are descriptive. The median row summarizes the ten task-specific coefficients.

Supplementary Table 3. BIOT against bipolar differential-entropy features

| Task | D, BIOT | D, bipolar DE | ΔS (points) | ΔU (points) | I (points) |
|---|---|---|---|---|---|
| FACED-9 | 0.22 | 0.49 | −13.0 [−15.8, −10.1] | −1.7 [−4.1, 0.5] | −11.4 [−13.9, −8.7] |
| FACED-2 | 0.15 | 0.36 | −5.5 [−9.9, −1.1] | −1.6 [−6.8, 3.7] | −3.9 [−7.5, −0.5] |
| SEED | 0.19 | 0.36 | −7.7 [−13.6, −2.2] | −6.1 [−12.6, −0.2] | −1.6 [−4.3, 1.0] |
| SEED-IV | 0.08 | 0.11 | −12.6 [−18.5, −6.8] | −12.3 [−18.7, −6.3] | −0.3 [−2.4, 1.8] |
| SEED-V | 0.09 | 0.22 | −9.6 [−16.1, −3.3] | −5.7 [−11.8, 0.3] | −3.8 [−7.7, 0.1] |
| SEED-VII | 0.11 | 0.24 | −8.6 [−12.3, −5.1] | −6.6 [−10.1, −3.1] | −2.0 [−4.5, 0.4] |
| DEAP-4 | 0.07 | 0.04 | 0.9 [−3.7, 5.3] | −1.2 [−5.8, 3.4] | 2.1 [−0.5, 4.7] |
| DEAP-2 | 0.05 | 0.04 | 1.6 [−3.8, 6.7] | 1.8 [−4.0, 7.5] | −0.2 [−2.8, 2.4] |
| DREAMER-9 | 0.14 | 0.31 | −5.6 [−11.2, −0.7] | 3.2 [−1.6, 8.1] | −8.8 [−16.5, −1.4] |
| DREAMER-2 | 0.10 | 0.22 | −4.5 [−13.2, 3.6] | 0.7 [−8.0, 9.6] | −5.3 [−18.0, 8.0] |

BIOT and bipolar differential-entropy (DE) features use the same 16 bipolar derivations. D is item identifiability. ΔS and ΔU are the advantages of BIOT under shared-material and new-material evaluation, respectively; I = ΔS − ΔU. Accuracy differences are in percentage points. Values summarize three seeds, and brackets give 95% confidence intervals from 2,000 crossed-bootstrap draws over participants and items.

The paired values accompany the method comparisons in the main text and Extended Data Fig. 6. BIOT had lower item identifiability than bipolar DE in eight of ten tasks, and a smaller shared-material gain in every task where the interaction interval excluded zero.

Supplementary Table 4. Calibration of the crossed bootstrap

| Task | Coverage: zero gain | Coverage: zero gain, item-heavy | Coverage: observed gain | Coverage: observed gain, item-heavy | Rejection: zero gain | Rejection: zero gain, item-heavy |
|---|---|---|---|---|---|---|
| FACED-9 | 0.917 | 0.906 | 0.927 | 0.901 | 0.064 | 0.077 |
| FACED-2 | 0.970 | 0.950 | 0.964 | 0.940 | 0.036 | 0.047 |
| SEED | 0.942 | 0.908 | 0.931 | 0.899 | 0.065 | 0.079 |
| SEED-IV | 0.996 | 0.984 | 0.995 | 0.987 | 0.009 | 0.018 |
| SEED-V | 0.988 | 0.985 | 0.988 | 0.983 | 0.014 | 0.023 |
| SEED-VII | 0.979 | 0.970 | 0.985 | 0.977 | 0.022 | 0.032 |
| DEAP-4 | 0.996 | 0.968 | 0.995 | 0.968 | 0.004 | 0.041 |
| DEAP-2 | 0.999 | 0.979 | 0.997 | 0.970 | 0.006 | 0.035 |
| DREAMER-9 | 0.955 | 0.954 | 0.958 | 0.959 | 0.042 | 0.050 |
| DREAMER-2 | 0.981 | 0.976 | 0.978 | 0.970 | 0.030 | 0.028 |

Empirical coverage of the nominal 95% percentile interval and rejection rate of the one-sided test at $\alpha = 0.05$, for differential-entropy features. Each scenario used 2,000 simulated datasets and 1,999 bootstrap replicates per dataset, as described in Supplementary Note 2. Item-heavy scenarios set the item standard deviation to max(1.5 × estimate, 6 percentage points). Values are proportions rounded to three decimal places; the item-heavy scenario with the observed gain also retained the empirical skew of the item effects.
